# Addressing Inconsistency in Functional Neuroimaging: A Replicable Data-Driven Multi-Scale Functional Atlas for Canonical Brain Networks

**DOI:** 10.1101/2024.09.09.612129

**Authors:** Kyle M. Jensen, Jessica A. Turner, Lucina Q. Uddin, Vince D. Calhoun, Armin Iraji

## Abstract

The advent of multiple neuroimaging methodologies has greatly aided in the conceptualization of large-scale functional brain networks in the field of cognitive neuroscience. However, there is inconsistency across studies in both nomenclature and the functional entities being described. There is a need for a unifying framework that standardizes terminology across studies while also bringing analyses and results into the same reference space. Here we present a whole-brain atlas of canonical functional brain networks derived from more than 100,000 resting-state fMRI datasets. These data-driven functional networks are highly replicable across datasets and capture information from multiple spatial scales. We have organized, labeled, and described the networks with terms familiar to the fields of cognitive and affective neuroscience in order to optimize their utility in future neuroimaging analyses and enhance the accessibility of new findings. The benefits of this atlas are not limited to future template-based or reference-guided analyses, but also extend to other data-driven neuroimaging approaches across modalities, such as those using blind independent component analysis (ICA). Future studies utilizing this atlas will contribute to greater harmonization and standardization in functional neuroimaging research.

## 1. Addressing Inconsistency in Neuroscience

### 1.1. Inconsistent Nomenclature: The Need for Standardized Terminology

The field of cognitive neuroscience has widely adopted the view that cognition is supported by several large-scale functional networks in the brain (Bressler & Menon, 2010). This paradigm has inspired many efforts to study underlying mechanisms of cognition by mapping and recording activity in the brain through neuroimaging and by applying various analytical approaches to the data produced (Bassett & Sporns, 2017). However, the field has yet to reach a consensus on basic terminology with regards to large-scale functional brain networks (Uddin et al., 2019). This presents a significant challenge to progress in the field, as inconsistency in labeling and terminology impedes the discoverability and propagation of researchers’ findings (Uddin et al., 2019, 2023; Winston, 2018), which introduces bias as one study may be favored over another merely due to a variation in terminology. Not only can it be ineffective for researchers studying the same regions and networks in the brain to describe them with different terminology, but it can also be inefficient as it leads to miscommunication and confusion between researchers within the field, students and trainees being initiated into the field, and potential benefactors of the research produced by the field (Uddin et al., 2019, 2023). This issue is particularly pernicious when it is extended to clinical neuroscience, where such miscommunication can hinder progress in the development of biomarkers, monitoring of disease progression and response to treatment, and identification of targets for neuromodulation. Indeed, the impact of inconsistent network nomenclature is far-reaching. As the field of neuroscience strives to achieve a consensus on key terminology, progress will help to unite related research findings and the field will develop a more comprehensive and complete understanding of the brain regions and networks under study.

### 1.2. Inconsistencies in the Application of Terminology: The Need for a Universal Reference Space

Another barrier to progress in the field occurs when neuroscientists use the same term to refer to different entities (Uddin et al., 2023). This may manifest as multiple studies using the same term (e.g., salience network) to describe different brain regions (Kong et al., 2024). This point is demonstrated very effectively by Kong and colleagues (2024) who illustrate an example where the anatomical regions labeled as the “salience” network across multiple popular brain atlases are largely non-overlapping in their spatial topographies. Extending upon this issue, there is a great amount of variability introduced by differences in methodology across studies. These include differences in regions of interest (ROIs; e.g., changes in seed location; M.-T. Li et al., 2023; Yeo et al., 2011), datasets (Elliott et al., 2019), and analytical approach, which all contribute to differences in observed functional patterns across studies. Furthermore, there is a great amount of variability across subjects in the same study (see Figure 1; Braga & Buckner, 2017), as well as within the same subject over the duration of the fMRI data acquisition (Iraji, Deramus, et al., 2019; Iraji, Fu, et al., 2019). Thus, a spatially fixed node, seed, or ROI may not represent the same functional unit from one participant to another due to individual differences, and it may not represent the same functional unit even within the same participant as sources vary spatially over time (Iraji et al., 2020). These particular challenges are not resolved by consistent nomenclature, instead, there is a need for a common set of functional units and data-driven methods which enable the consistent adoption of these units across subjects, datasets, and studies.

**Figure 1.**
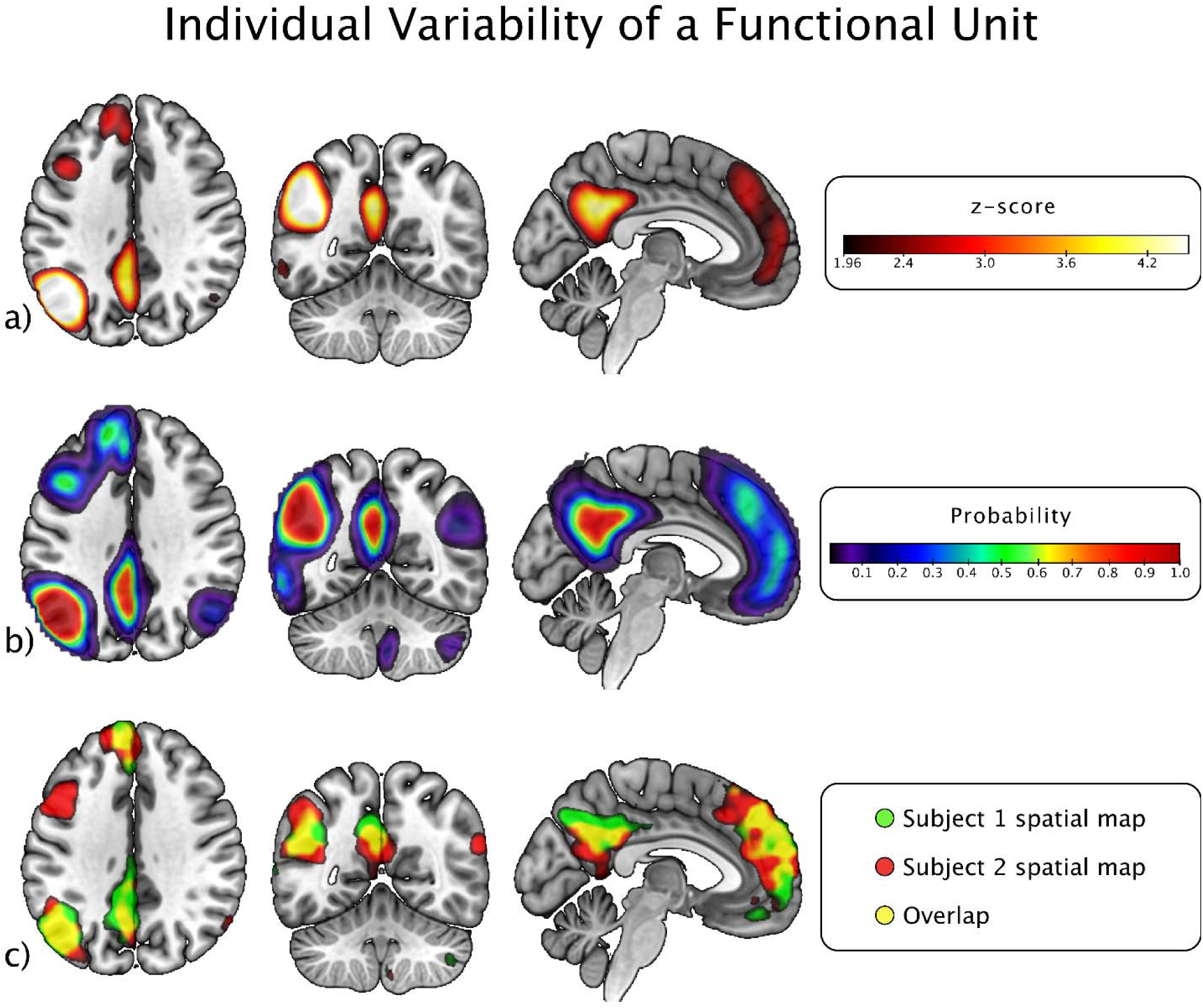
Individual variability of a functional unit. The first panel displays (a) the average of individual subject spatial maps for an intrinsic connectivity network (ICN) extracted from ∼40k subjects in the UK Biobank dataset overlayed on the MNI152 brain template. All voxels represent z-scores and only significant (z-score > 1.96) voxels are displayed. For each subject we calculated (b) the probability that a given voxel of the selected ICN would be significant (z-score > 1.96) across all ∼40k subjects from the UK Biobank dataset. This probability spatial map is overlayed on the MNI152 brain template using the NIH color scale, where color represents probability (e.g., purple is a probability of ∼10%, green is ∼50%, red is around the range of 80-100%). This probability map has a lower threshold of 1%. For panel (c) spatial maps for two subjects are plotted in green and red overlayed on the MNI152 brain template. The color is additive, with yellow representing overlap between the spatial maps of the two subjects. There is relatively little overlap, demonstrating a great amount of individual variability in the spatial map of the given ICN. The voxels in these spatial maps are z-scored, with only significant (> 1.96) voxels displayed.

### 1.3. The Power and Limitations of an Atlas: Lessons from Brodmann

More than a century ago, the German neurologist Korbinian Brodmann defined 52 distinct brain regions (43 in humans) based on cellular structural characteristics (i.e., cytoarchitecture; Brodmann, 1909; for English translation see Brodmann & Gary, 2006). Brodmann’s work established a referential atlas which is still widely used today (Zilles & Amunts, 2010). The Brodmann Areas (BAs) have great utility because they help link different studies to objectively^1^ defined universally known anatomical regions, even when there may be great variability in the functionally and anatomically descriptive names for a given region within the field of neuroscience. However, even this taxonomy falls short in unifying the field of neuroscience, as it is not informed by modern neuroimaging techniques and is limited to describing anatomy (i.e., it is not well equipped to describe functional units), it does not describe the whole brain (i.e., it omits subcortical and cerebellar structures), and it is not sensitive to individual differences (Zilles & Amunts, 2010). Even a century later, there remains a great need for a common reference space for the functional brain which is adaptable across studies and individual participants within them. Modern data-driven approaches have demonstrated great promise in overcoming these challenges (Iraji, Fu, et al., 2019; Iraji et al., 2020; Luo et al., 2021; Wang et al., 2015).

### 1.4. NeuroMark: A Data-Driven Approach Towards a Common Reference Space

NeuroMark is a term coined by Du and colleagues (2020) which represents an ongoing effort toward establishing a common reference space across functional imaging analyses in the field of neuroscience. Past efforts (Du et al., 2020) have included the development of a standardized fully automated preprocessing and analysis pipeline which incorporates a template used as a reference for spatially constrained independent component analysis (ICA; Calhoun et al., 2001, 2009) of functional magnetic resonance imaging (fMRI) data. The functionally distinct entities or units extracted through ICA represent coherent brain activity and are referred to as intrinsic connectivity networks (ICNs). The NeuroMark_fMRI_1.0 template includes 53 highly replicable ICNs extracted from 1828 control subject resting-state fMRI scans and validated across 2442 clinical subject resting-state fMRI scans. This template is implemented in the Group ICA of fMRI Toolbox (GIFT; http://trendscenter.org/software/gift; and separately available at http://trendscenter.org/data; Iraji et al., 2021) and has been successfully utilized by many prior studies to identify and describe unique patterns of brain connectivity associated with different clinical groups (Du et al., 2020; Fu, Iraji, Sui, et al., 2021; Fu, Iraji, Turner, et al., 2021; Fu, Sui, et al., 2021; Jensen et al., 2024; K. Li et al., 2021). This data-driven approach has many advantages which address the limitations of subject and dataset variability. In short, prior work suggests that ICA is better equipped than ROI-based approaches for extracting functional connectivity features retaining individual-level variability (Du et al., 2020; Yu et al., 2017) and group ICA methods overcome the limitations of a potential lack of spatial correspondence between subjects (Calhoun et al., 2001). One potential caveat of blind-ICA, like any pure data-driven approach, is that the extracted ICNs do not necessarily have a one-to-one correspondence between studies (Du et al., 2020; Iraji et al., 2023). To address this limitation, the NeuroMark pipeline implements an a priori-driven (i.e., spatially constrained) ICA which is informed by a reliable whole-brain network template to overcome inconsistency in components across datasets and studies (Du et al., 2020; Q.-H. Lin et al., 2010). Together, these characteristics make the NeuroMark approach well-equipped for comparing findings across studies, as it is sensitive to individual differences but also brings those findings into the same reference space across datasets. Recently, Iraji and colleagues (2023) sought to further improve the reliability of the NeuroMark networks by utilizing a large-scale (N > 100,000) resting-state fMRI (rsfMRI) dataset to develop a multi-spatial-scale network template with 105 highly replicable ICNs covering the whole brain. In data-driven approaches utilizing ICA, different model orders (i.e., the number of estimated components) capture information at different spatial scales (Iraji et al., 2022; H. Li et al., 2018). Multi-model-order spatial ICA (msICA) captures scale specific complementary information across multiple spatial scales (Iraji et al., 2022). In other words, the resulting parcels or ICNs from msICA improve upon other ICA approaches by offering more reliable and informative coverage of the brain. Iraji and colleagues (2023) utilized msICA to identify networks which were consistent across multiple model orders, or spatial scales, ranging from 25 (i.e., the ICA estimates 25 independent components) to 200 for a total of 900 unique ICNs. Stringent criteria were applied (e.g., identifying the most stable ICNs across 100 iterations of ICA; see Iraji et al., 2023) to identify the most reliable ICNs, and only the most spatially distinct (spatial similarity < 0.8 with all other ICNs) ICNs were selected for the final template. Thus, the 105 ICNs in the resulting template represent unique sources of functional brain circuitry that are reliable across individual subjects across the lifespan, as well as multiple spatial scales (Bajracharya et al., 2024; Iraji et al., 2023).

While Iraji and colleagues (2023) identified 105 highly reliable and replicable multi-scale networks, they did not make an attempt to label, group, and describe them. Admittedly, this is a challenging task, which typically involves anatomical labels being imposed upon functional units that reflect unique patterns of connectivity. In other words, ICNs represent patterns of functional connectivity that often span across many anatomical features or boundaries that would typically be used as topographical markers. Due to their unique properties (e.g., their spatial topography and fluidity across subjects and over time), not only is it difficult to describe these functional nodes in anatomical terms, but in doing so we risk introducing bias and inaccuracy. Conversely, the lack of a standardized structure or framework for organizing the results would considerably increase the amount of time and effort required by investigators to interpret and convey their results and would introduce unnecessary variability and inconsistency in terminology, both of which would greatly undermine the utility of the NeuroMark approach. Therefore, we pose that the creation of such a framework is warranted as long as it is done with great transparency, is sufficiently descriptive, and is utilized appropriately. Additionally, the framework would likely benefit from updates over time.

### 1.5. Translating NeuroMark across the Field of Neuroscience

Here we sought to improve the accessibility of the NeuroMark multi-scale template by 1) assigning individual labels to each of the 105 ICNs which describe their spatial overlap with anatomical structures, 2) assigning domain and subdomain labels for groups of spatially similar ICNs which utilize terminology commonly used in neuroscience and neuroimaging research, and 3) re-ordering the ICNs to fit within these groups and to cluster them with other spatially similar ICNs within these groups. We anticipate that these changes will make the existing NeuroMark template more accessible and interpretable by other investigators within the field of neuroscience.

Each of the 105 ICNs was visually inspected in MRIcroGL (Rorden & Brett, 2000; available at https://www.nitrc.org/projects/mricrogl) overlayed on the MNI152 template (Mazziotta et al., 2001). Notes were taken on the location of the peak coordinates as well as overall size and shape of the ICN. Each ICN was viewed in overlay with multiple different atlases including Brodmann Areas (BAs; atlas available in MRIcroGL; see Zilles & Amunts, 2010 for description and historical significance) and the Automated Anatomical Atlas (AAL; Tzourio-Mazoyer et al., 2002). In addition, for each of the previously mentioned atlases, we calculated the total number of significant voxels within each atlas region which overlapped with each ICN (see the chord charts in Figure 2 and Figure S3; for plotting chord charts see Liu, 2024) and identified the top ten regions (e.g., BAs or AAL ROIs) corresponding with each ICN. We also compared our assigned labels against cognitive and behavioral neuroscience textbooks and research articles, search engines including Neurosynth.org, Google Scholar, and Wikipedia, as well as common knowledge. The resulting domain, subdomain, and individual labels have been released with NeuroMark version 2.2 (freely available in GIFT: http://trendscenter.org/software/gift; Iraji et al., 2021; and separately available at http://trendscenter.org/data) and are presented in order in **Table 1**, with associated spatial maps displayed in composite view in **Figure 2** and individually in supplementary materials **Figure S1**.

**Figure 2.**
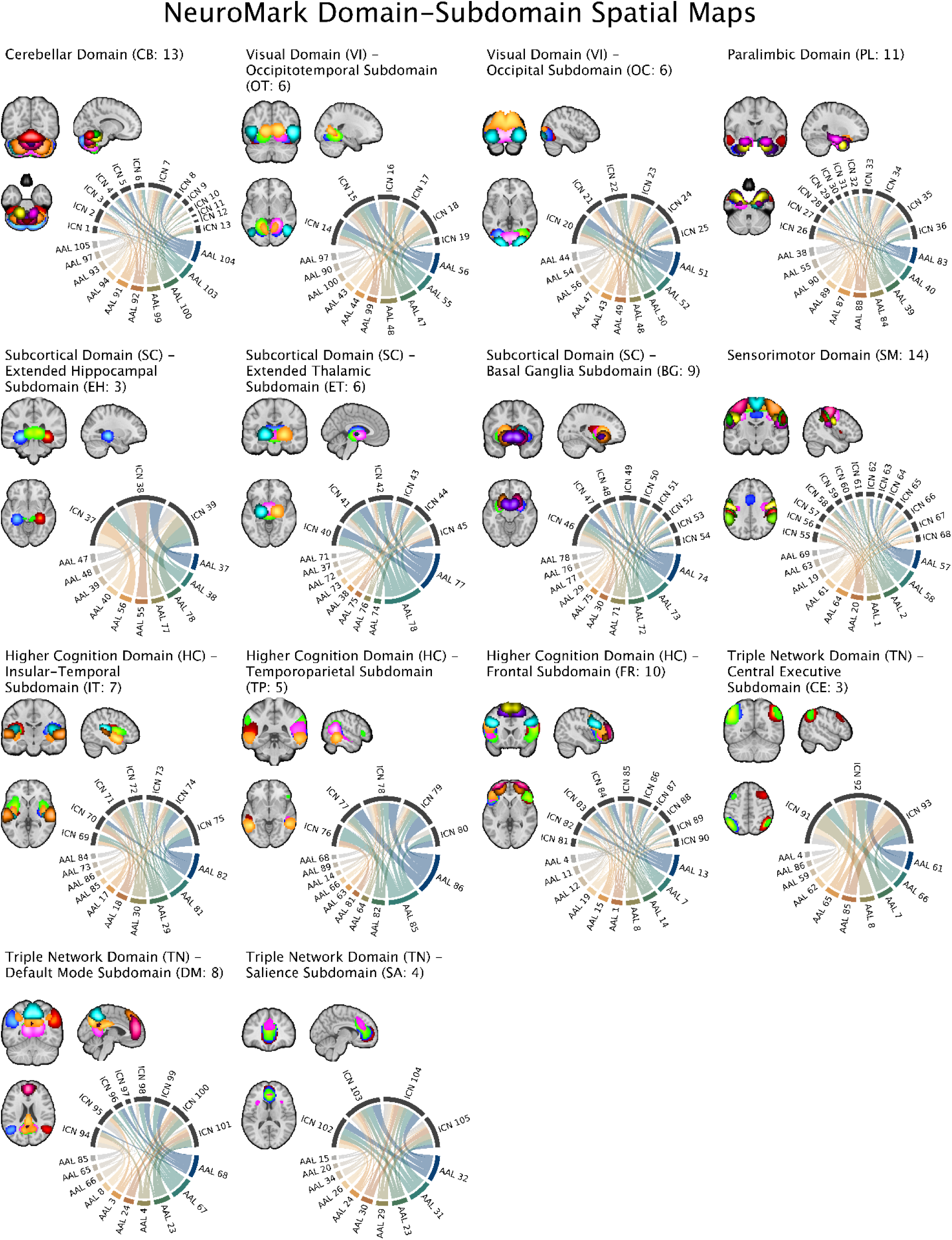
Overlayed spatial maps of the 105 intrinsic connectivity networks (ICNs) from the NeuroMark 2.2 multi-scale template are plotted above for each of the seven domains and 14 subdomains: cerebellar (CB), visual-occipitotemporal (VI-OT), visual-occipital (VI-OC), paralimbic (PL), subcortical-extended hippocampal (SC-EH), subcortical-extended thalamic (SC-ET), subcortical-basal ganglia (SC-BG), sensorimotor (SM), higher cognition-insular temporal (HC-IT), higher cognition-temporoparietal (HC-TP), higher cognition-frontal (HC-FR), triple network-central executive (TN-CE), triple network-default mode (TN-DM), and triple network-salience (TN-SA). All spatial maps have been thresholded to display ICN voxels with a z-score > 3. We used a higher threshold than *z* = 1.96 (*p* = .05) to make it easier to visualize and distinguish between the spatial maps of individual components. In addition, chord charts are plotted for each subdomain, displaying the proportion of significant (z-score > 1.96, p-value < 0.05) voxels overlapping with the top 10 most contributing regions from the automated anatomical atlas (AAL). A reference table for the AAL regions (see Table S2) as well as chord charts with the top 10 Brodmann Areas overlapping with each subdomain (see Figure S3) are provided in the supplementary materials.

**Table 1.**
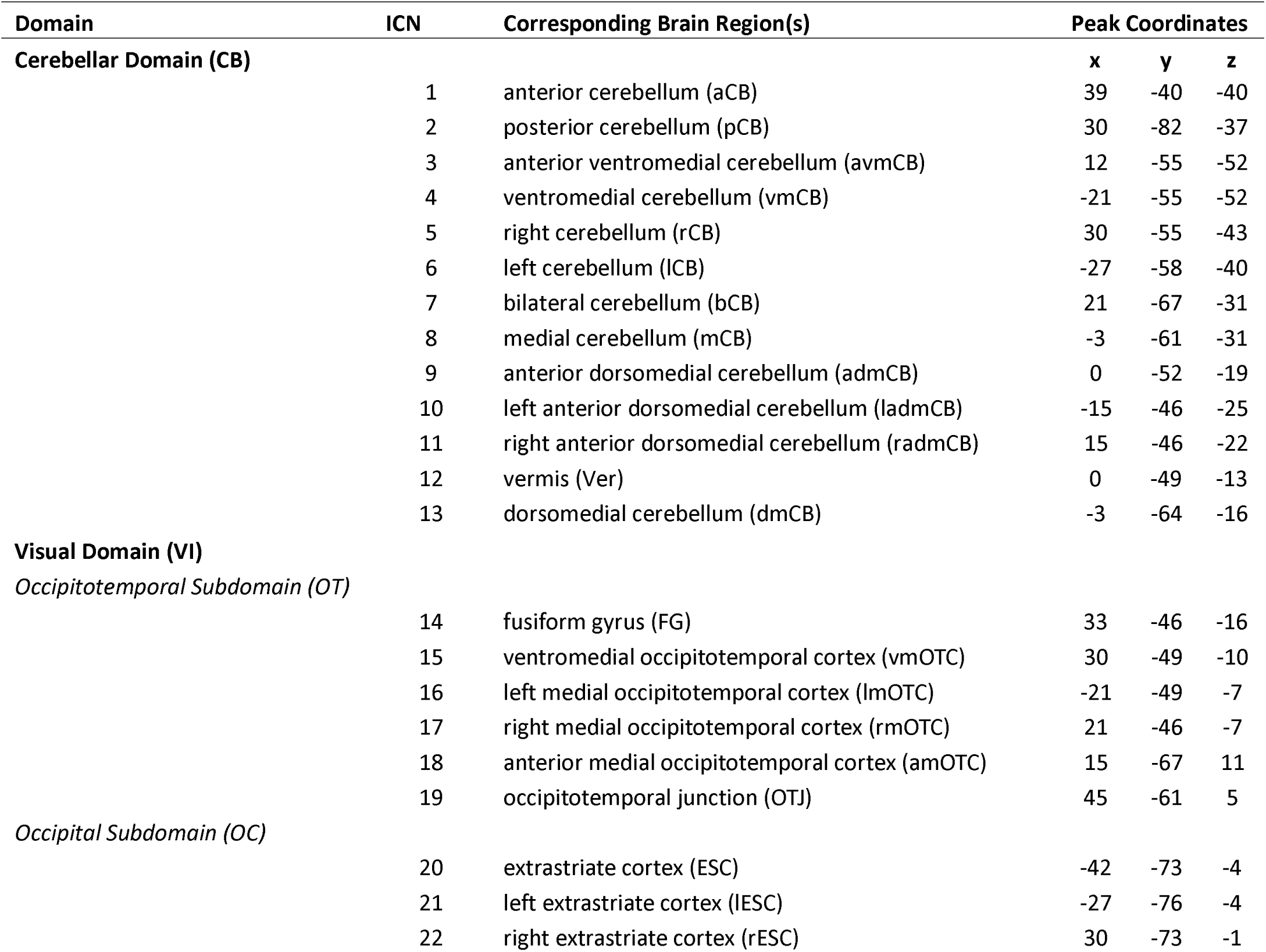
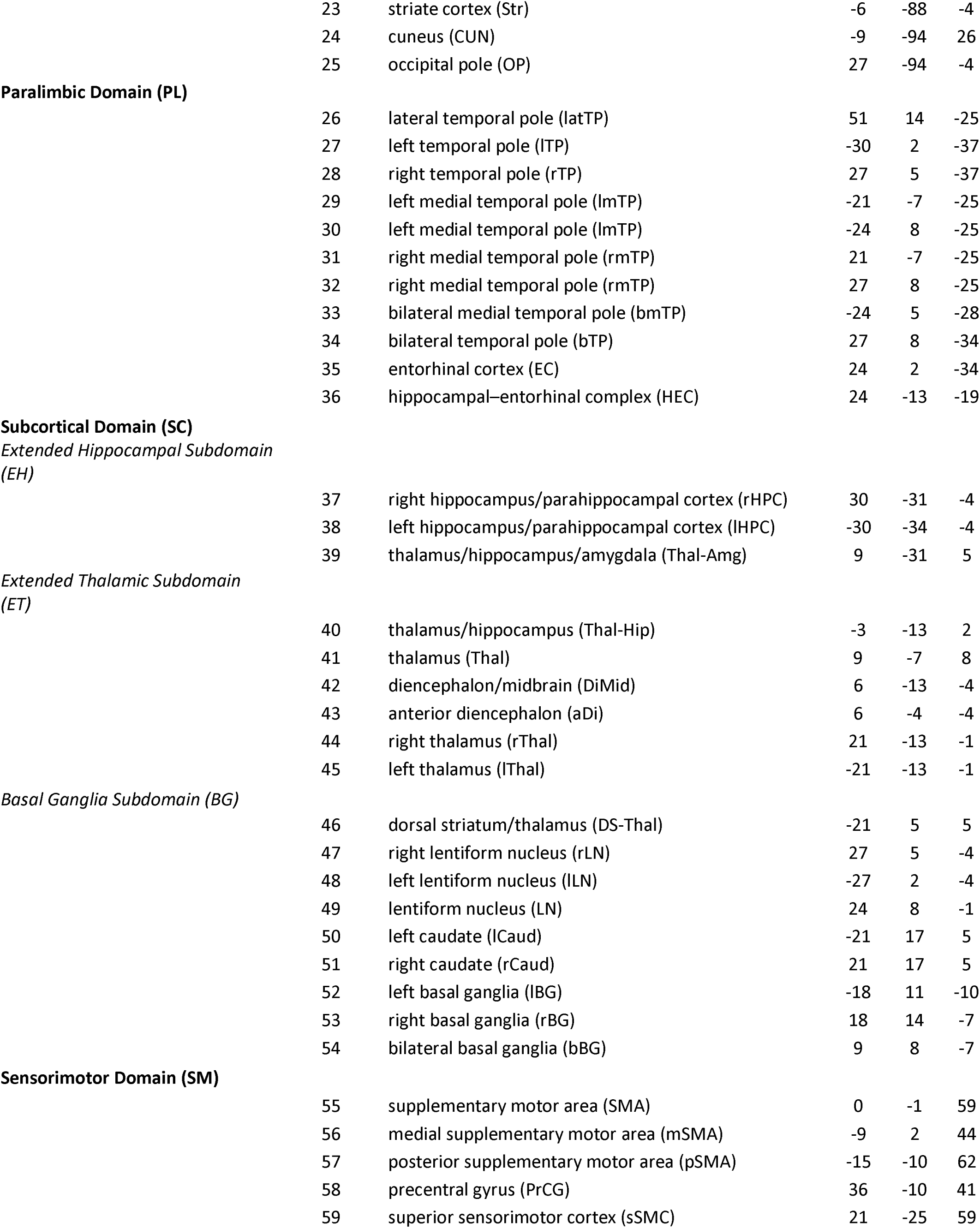
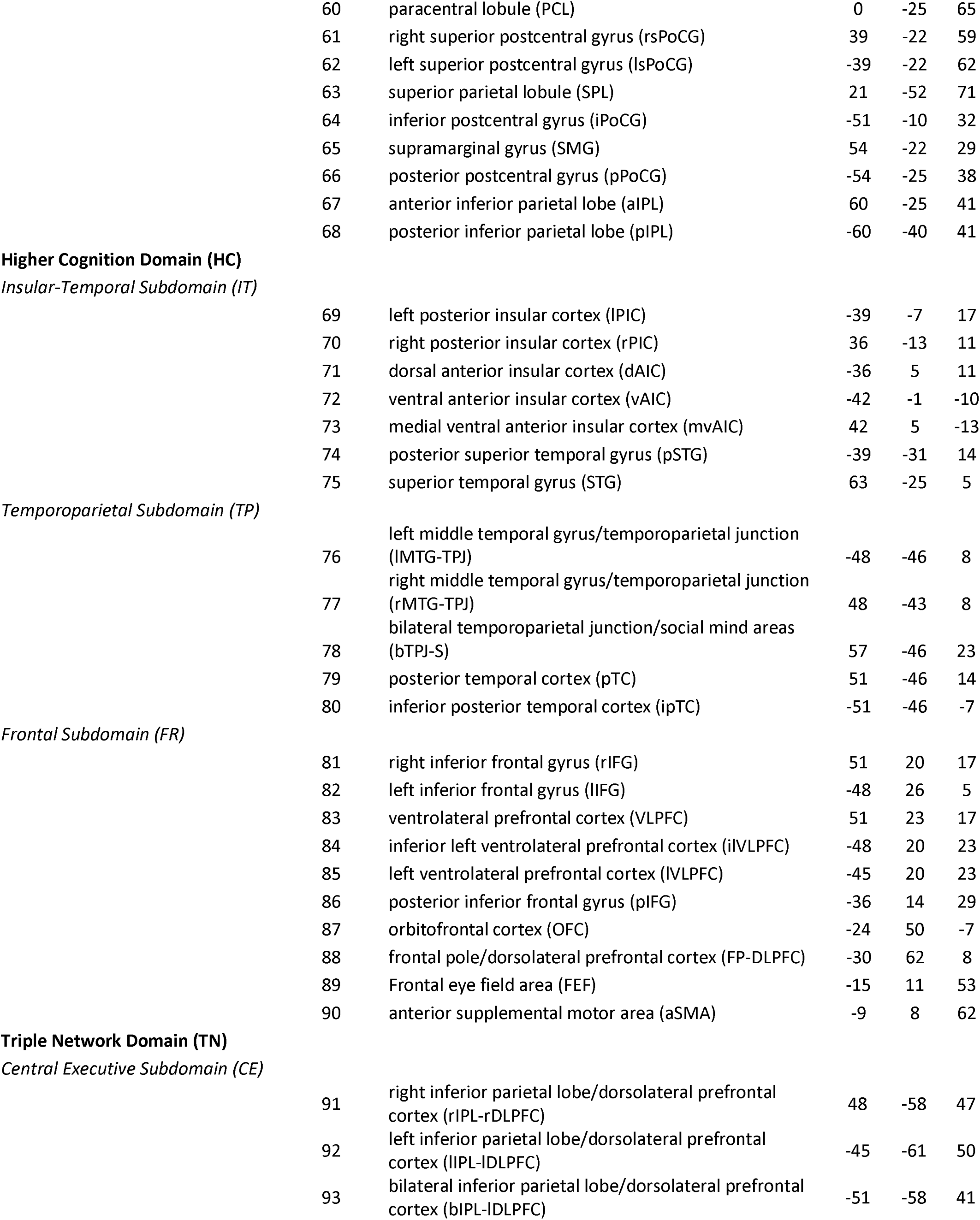
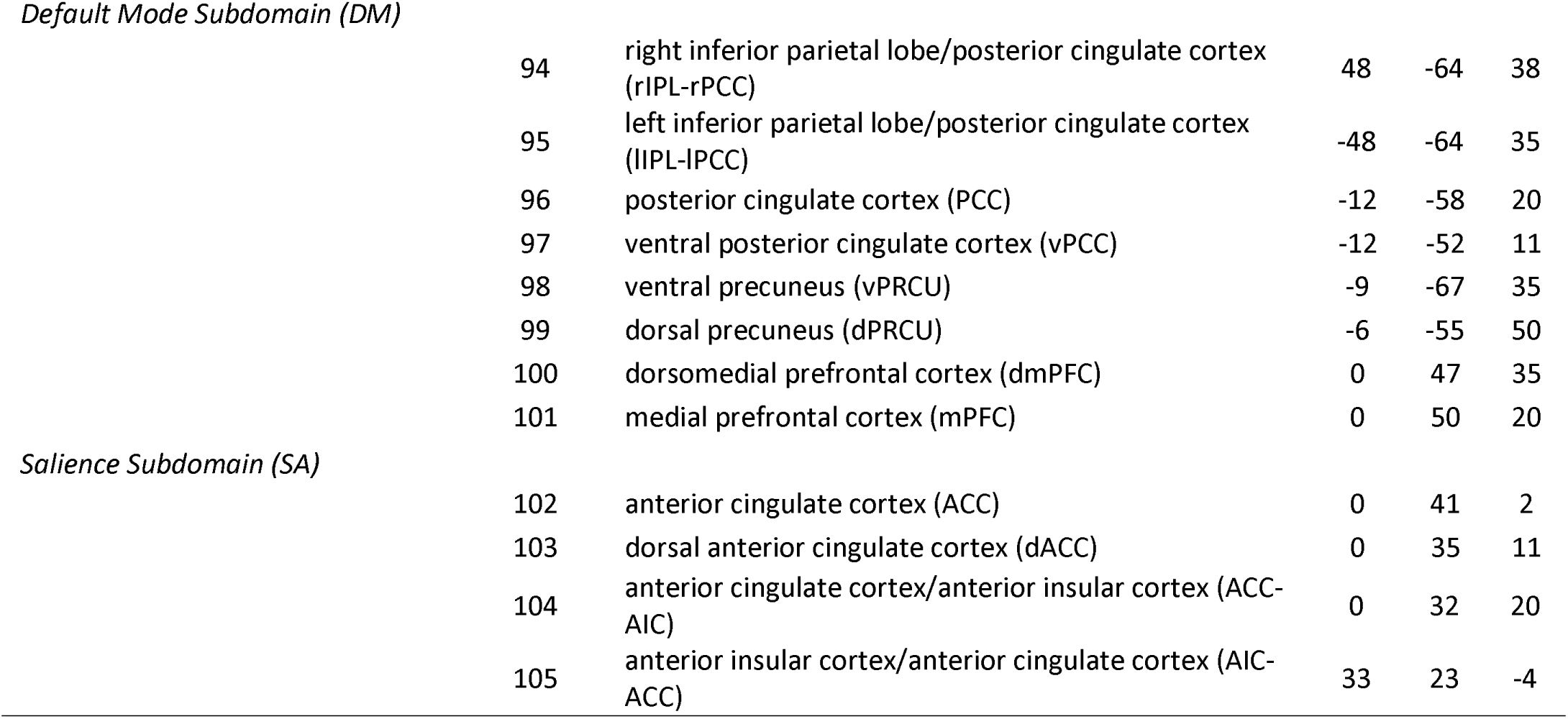
The 105 intrinsic connectivity networks (ICNs) from the NeuroMark 2.2 multi-scale template are grouped into seven domains and 14 subdomains based on spatially overlapping anatomical or functionally defined brain regions: cerebellar (CB), visual-occipitotemporal (VI-OT), visual-occipital (VI-OC), paralimbic (PL), subcortical-extended hippocampal (SC-EH), subcortical-extended thalamic (SC-ET), subcortical-basal ganglia (SC-BG), sensorimotor (SM), higher cognition-insular temporal (HC-IT), higher cognition-temporoparietal (HC-TP), higher cognition-frontal (HC-FR), triple network-central executive (TN-CE), triple network-default mode (TN-DM), and triple network-salience (TN-SA). The MNI space coordinates (x,y,z) for peak location are displayed to the right of each ICN.

Previous studies have used subject-specific ICNs such as these to draw insight from brain imaging data (Du et al., 2020; Iraji et al., 2023). Specifically, the relationship between the extracted ICNs is typically referred to as functional network connectivity (FNC) and can be represented with a Pearson correlation between the time courses of two ICNs. Consequently, ICNs which are functionally related are strongly correlated or anticorrelated (represented with a higher positive or negative coefficient *r*). Similarly, ICNs which are spatially similar tend to also have functional overlap, which manifests as stronger FNC. When FNC is plotted in a matrix the correlations between related ICNs form distinct patterns of modularity. Thus, insight into the relationship between activity in different brain regions can be drawn by examining an FNC matrix. Since the NeuroMark approach spatially constrains the extracted ICNs to a template, we would expect to observe similar patterns in FNC from one dataset to the next. To demonstrate this, we plotted the mean of subject-specific FNC across 39,342 subjects from the UK Biobank dataset (Littlejohns et al., 2020; see Figure 3a), 1,010 subjects from the HCP dataset (Van Essen et al., 2013; see Figure 3b), and 11,426 subjects from the ABCD dataset (Casey et al., 2018; see Figure 3c), all of which are large and widely used resting-state fMRI datasets. All three datasets^2^ displayed highly similar patterns of FNC, demonstrating the effectiveness of the NeuroMark approach in bringing functional data from different subjects and datasets into the same reference space.

**Figure 3.**
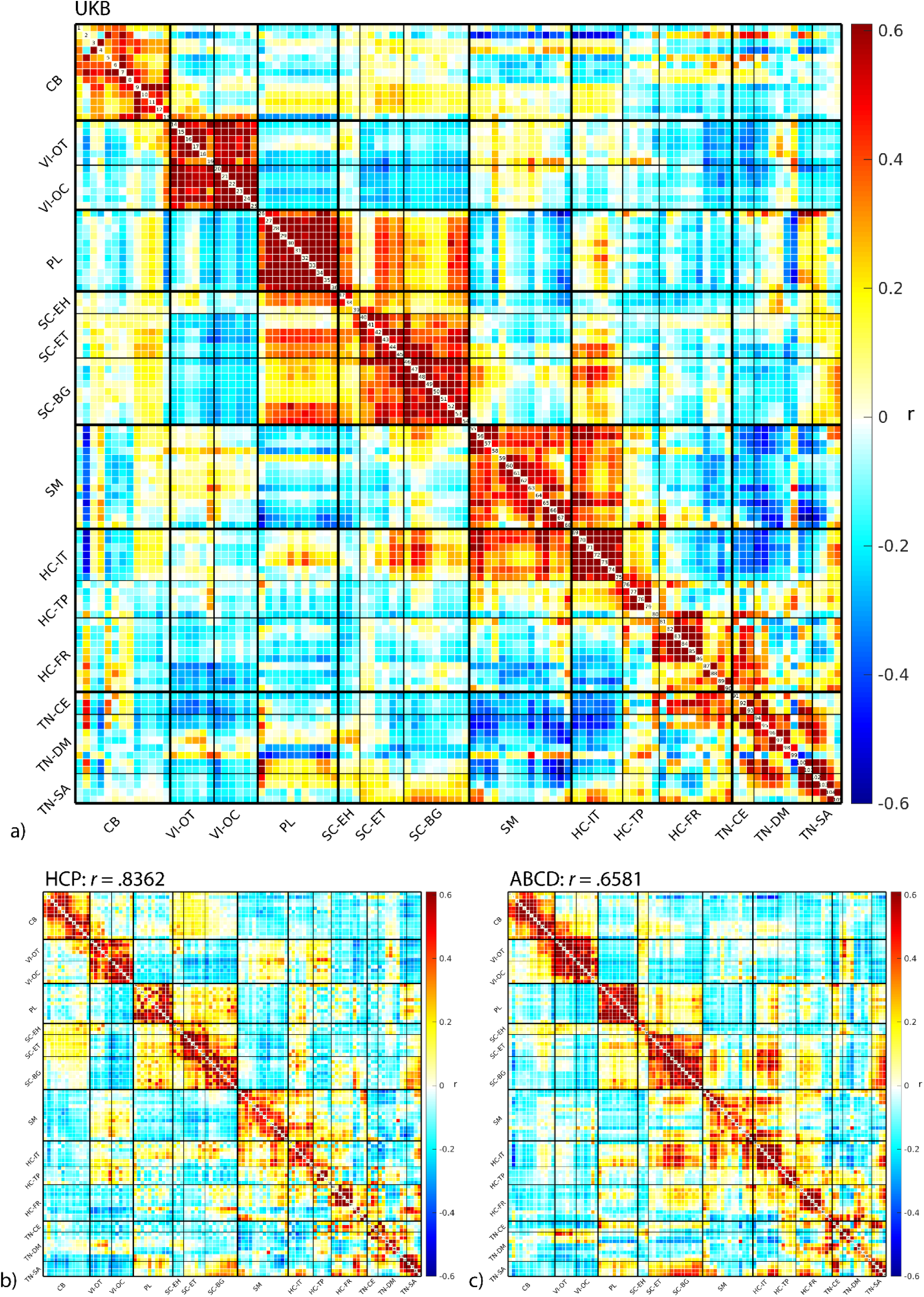
Functional network connectivity (FNC) averaged across (a) 39,342 participants from the UKB dataset is displayed in the matrix above with red indicating higher positive values (Pearson correlation coefficient *r*) and blue indicating lower negative values. Intrinsic connectivity networks (ICNs) are grouped by their domain-subdomain labels, consistent with the NeuroMark_fMRI_2.2 multi-scale template: cerebellar (CB), visual-occipitotemporal (VI-OT), visual-occipital (VI-OC), paralimbic (PL), subcortical-extended hippocampal (SC-EH), subcortical-extended thalamic (SC-ET), subcortical-basal ganglia (SC-BG), sensorimotor (SM), higher cognition-insular temporal (HC-IT), higher cognition-temporoparietal (HC-TP), higher cognition-frontal (HC-FR), triple network-central executive (TN-CE), triple network-default mode (TN-DM), and triple network-salience (TN-SA). The reliability of the this template is demonstrated by the high similarity (Pearson correlation) between the UKB FNC and the mean FNC for (b) 1,010 participants from the HCP dataset and (c) 11,426 participants from the ABCD dataset.

## 2. The NeuroMark Domains

The following sections are intended to provide a brief definition and description for each domain and subdomain in the NeuroMark atlas. While some notable characteristics and functions are attributed to the brain structures found within each section, the sections below are far from comprehensive and are intended to supplement and offer direction in the interpretation of the results of future analyses. Caution should be used to avoid restricting or replacing more thorough consideration, which risks introducing a bias in the interpretation of results.

### 2.1. Cerebellar Domain

The cerebellar domain (CB) includes 13 ICNs (see ICNs 1-13), all within the cerebellum. The cerebellum is a large brain structure which is often overlooked in cognitive and affective neuroscience and has been omitted from many brain atlases, however, the cerebellum is gaining increased attention in rsfMRI studies (Buckner et al., 2011; Habas, 2021; Iraji, Deramus, et al., 2019) and has been incorporated in models for psychiatric illness (Andreasen et al., 1998; Harikumar et al., 2023; Pinheiro et al., 2021; Rudolph et al., 2023). This is likely because sensorimotor control was historically considered the primary function of the cerebellum, although, modern views also recognize its involvement in cognitive, affective, and social processing (Buckner, 2013; Habas, 2021; Rudolph et al., 2023; Van Overwalle et al., 2020). In particular, emotional processes have been attributed to the vermis (see ICNs 8-13; Ciapponi et al., 2023; Habas, 2021) a medial structure dividing the two lobes of the cerebellum, and social processes have been attributed to posterior regions of the cerebellum (see ICN 2; Van Overwalle et al., 2020). Notably, like other brain regions such as the insula (Uddin et al., 2011), the cerebellum’s involvement with large-scale networks is complex and appears to change over time (i.e., across development; Clark et al., 2022). However, investigations in spatial dynamics have specifically demonstrated that cerebellar contributions emerge at particular timepoints or states, which can make it potentially difficult to capture the dynamic role of the cerebellum through static analyses (Iraji, Deramus, et al., 2019). It should also be noted that many reference books and articles classify the cerebellum as a subcortical structure, however, we have placed the cerebellum in its own domain to help delineate its unique functional contributions and distinguish it from other subcortical structures.

### 2.2. Visual Domain

The visual domain (VI) contains 12 ICNs (see ICNs 14-25), which have been divided into two subdomains, the occipitotemporal subdomain (OT) and the occipital subdomain (OC). These regions of the brain are highly specialized, contributing primarily to the processing of visual information (Huff et al., 2024).

#### 2.2.1. Visual Domain - Occipitotemporal Subdomain

The OT includes six ICNs (see ICNs 14-19) in occipital and temporal regions. Structures in the inferior temporal lobe, particularly the fusiform gyrus (BA 37; see ICNs 14-17), are known to perform various roles in high-level visual processing including object recognition, face perception, reading, and memory (Weiner & Zilles, 2016). Indeed, the inferior temporal lobe is a key structure in the ventral visual stream, also known as the “what” pathway for visual processing (Goodale & Milner, 1992).

#### 2.2.2. Visual Domain - Occipital Subdomain

The OC includes six ICNs (see ICNs 20-25) centered in the striate cortex (corresponding with BA 17; see ICN 23), which is functionally known as the primary visual cortex or visual area 1 (V1), and the extrastriate cortex (corresponding with BA 18 and BA 19; see ICNs 20-22), functionally referred to as visual association areas or visual areas 2, 3, 4, and 5 (V2, V3, V4, and V5; Cotman & McGaugh, 1980; Huff et al., 2024). The primary visual cortex receives sensory input from the retina via the thalamus and serves as the lowest and most basic level of visual processing, with higher and more complex levels of processing occurring in visual association areas, extending to other regions of the cortex (Cotman & McGaugh, 1980; Huff et al., 2024).

### 2.3. Paralimbic Domain

The paralimbic domain (PL) includes 11 ICNs (see ICNs 26-36) centered in regions surrounding limbic structures (e.g., the hippocampus and amygdala), namely the entorhinal cortex (BA 28 and 34; see ICNs 35-36), medial temporal lobe (BA 35 and 36; see ICNs 29-33), and temporal pole (BA 38; see ICNs 26-28, 30, 32-34). Although it may not be commonly used, the term selected for this domain has been used previously to describe these structures (Juárez et al., 2013; Kiehl, 2006; Laurens et al., 2005; Mesulam, 2000), and it appears to describe this unique group of ICNs well, distinguishing them from the other domains presented in this atlas. Paralimbic structures contribute to perceptual processes, including gustation, olfaction, and the perception of pain and time, as well as a wide range of higher cognitive functions, including emotion, memory, language, and learning (Brown & Eldridge, 2008; Cleland & Linster, 2019; Herlin et al., 2021; Insel & Takehara-Nishiuchi, 2013; Mesulam, 2000).

### 2.4. Subcortical Domain

The subcortical domain (SC) includes 18 ICNs (see ICNs 37-54) which have been divided into three subdomains, the extended hippocampal subdomain (EH), the extended thalamic subdomain (ET), and the basal ganglia subdomain (BG). Along with the cerebellum, subcortical structures are excluded from many brain atlases and there is much room to improve our understanding of their functional contributions to higher cognition (Janacsek et al., 2022; Parvizi, 2009; Saban & Gabay, 2023). However, subcortical regions play a critical role in a vast array of cognitive processes (Janacsek et al., 2022) and are a key component of many psychiatric disease models (Andreasen et al., 1998; Harikumar et al., 2023). Although the ICNs in the current atlas do not map onto specific anatomical structures with great precision, due at least in part to the dense circuitry and relatively small size of subcortical structures (see Hollander et al., 2015), the unique shape and position of each ICN is informative and can be described and categorized into the following subdomains.

#### 2.4.1. Subcortical Domain - Extended Hippocampal Subdomain

The EH includes three ICNs (see ICNs 37-39) centered primarily in the hippocampus, but also extending into surrounding regions, namely the amygdala, thalamus, and parahippocampal cortex. These structures are central to the limbic system and are involved in explicit memory as well as various other functions, including emotion (e.g., fear and anxiety) and motivation, navigation, creativity, and imagination (Comrie et al., 2022; Lisman et al., 2017; Torrico & Abdijadid, 2024). The hippocampus in particular has been heavily studied in memory research and has been established as a central structure in the formation of long-term memory (Bird & Burgess, 2008; van Strien et al., 2009). However, modern views have emphasized that the hippocampus does not act alone, but rather it integrates and binds information from other brain structures and networks via its diverse neural circuitry (Borne et al., 2023; Whittington et al., 2022).

#### 2.4.2. Subcortical Domain - Extended Thalamic Subdomain

The ET includes six ICNs (see ICNs 40-45) which are centered in the thalamus, but extend into other structures of the diencephalon as well as the hippocampus and midbrain. The thalamus is located near the center of the brain and represents a major subdivision of the subcortex. The thalamus contains several nuclei which are intricately connected with other subcortical and cortical structures through extensive neural circuitry (Hwang et al., 2017). These nuclei are well known for their unique roles in limbic functions (i.e., emotion and motivation) and regulating sensory information (e.g., visual, auditory, pain) as well as arousal (i.e., sleep and wakefulness; Torrico & Munakomi, 2024). Although it has traditionally been viewed as a passive relay center, the thalamus is now understood to be actively involved in regulating information transfer between cortical regions (Hwang et al., 2017), playing a key role in many cognitive functions such as behavioral flexibility, language, and memory (Saalmann & Kastner, 2015).

#### 2.4.3. Subcortical Domain - Basal Ganglia Subdomain

The BG includes 9 ICNs (see ICNs 46-54) which are centered in the basal ganglia and overlap with parts of the thalamus. The largest structure in the basal ganglia is the striatum which consists of the caudate and lentiform nuclei (i.e., the putamen and globus pallidus), the substantia nigra, and the subthalamic nucleus (Utter & Basso, 2008; Young et al., 2024). The basal ganglia has strong connections with the sensorimotor cortex, thalamus, and brainstem, and plays a critical role in regulating voluntary motor movements, and is also involved in habit formation, learning, emotion, and decision-making (Jansson-Boyd & Bright, 2024; Nagano-Saito et al., 2014; Yin & Knowlton, 2006; Young et al., 2024).

### 2.5. Sensorimotor Domain

The sensorimotor domain (SM) includes 14 ICNs (see ICNs 55-68) and encompasses several structures in the frontal and parietal cortex including supplementary motor and premotor cortex, precentral and postcentral gyri, the paracentral lobule, and the supramarginal gyrus. The supplementary motor area (SMA; see ICNs 55-57) is a dorsomedial region of the frontal cortex (and superior subdivision of BA 6) and is anterior to the precentral gyrus. The SMA is heavily connected with other motor cortex and is specifically involved in planning motor movement (Tanji & Shima, 1994), as well as temporal sequencing and the perception of time duration (Cona & Semenza, 2017; Coull et al., 2016; Tanji & Shima, 1996). The premotor cortex is ventral to the SMA (the inferior subdivision of BA 6) and is also involved in planning and coordinating complex movement sequences, although while the SMA is related to internally-triggered movements (e.g., internally visualizing or thinking about the movement), the premotor cortex is related to externally-guided movements (e.g., incorporating visual feedback; Debaere et al., 2003; Sira & Mateer, 2014). The precentral gyrus (see ICN 58) forms the posterior border of the frontal cortex and contains the primary motor cortex (also known as area M1 or BA 4) which directly encodes neural signals to elicit movement via lower motor neurons in the brainstem and spinal cord (Purves et al., 2001b; Sira & Mateer, 2014). The postcentral gyrus (see ICNs 61-64, & 66) makes up the anterior border of the parietal lobe and contains the primary somatosensory cortex (also known as area S1 or BAs 1-3) which is the central brain region responsible for our perception of pressure, pain, touch, and temperature (Konstantopoulos & Giakoumettis, 2023; Raju & Tadi, 2024; Treede & Apkarian, 2008). Notably, the parietal lobe processes other types of sensory information as well. The dorsal visual and auditory streams lead from the primary visual (V1) and auditory (A1) cortex to the parietal lobe, forming the “where” pathway for processing spatial visual and auditory information (Goodale & Milner, 1992; Rauschecker & Tian, 2000). The paracentral lobule (see ICN 60) forms the medial surface of the precentral and postcentral gyri and, like the lateral surface, is associated with sensory and motor functions, particularly relating to the lower half of the body (Johns, 2014). The supramarginal gyrus (BA 40; see ICNs 65 & 67) is located in the inferior parietal lobe (IPL) and contributes to language, specifically phonologic working memory and speech production and perception (Gow, 2012; Vigneau et al., 2006).

### 2.6. Higher Cognition Domain

The higher cognition domain (HC) is the largest domain, encompassing most of the cerebral cortex. This domain consists of 22 ICNs (see ICNs 69-90) which have been divided into 3 subdomains, the insular-temporal subdomain (IT), the temporoparietal subdomain (TP), and the frontal subdomain (FR). This domain represents a group of brain regions which are highly interconnected with other domains, as demonstrated in the modularity of the FNC matrix (see Figure 3), and have been studied more extensively than other brain regions in their involvement in complex brain functions such as language and communication, problem-solving, math, decision-making, social and moral judgment, and imagination and creativity (Saban & Gabay, 2023; Schall, 2009). Consequently, these regions of the brain may also be the most difficult to categorize and compartmentalize into a single domain due to their integrated nature and the large range of cognitive functions associated with them.

#### 2.6.1. Higher Cognition Domain - Insular-Temporal Subdomain

The IT consists of seven ICNs (see ICNs 69-75) in the insular cortex and temporal lobe, particularly the superior temporal gyrus (STG; BA 41, 42, & 22). The insula (see ICNs 69-73) is an under-studied brain region located within the Sylvian fissure, between the IPL and temporal lobe. Functionally, the insula is extremely heterogenous, contributing to sensorimotor processing (e.g., visceral, somatic, auditory, chemosensory, etc.), socio-emotional processing (e.g., emotional experience, empathy, social cognition, etc.), attention, and language/speech (Uddin et al., 2017). Uddin (2015) highlights how these different functions might be attributed to specific subdivisions of the insular cortex, specifically, the posterior insular cortex (PIC) is believed to mediate sensorimotor processes, the dorsal anterior insular cortex (dAIC) is believed to mediate cognitive processes, and the ventral anterior insular cortex (vAIC) is believed to mediate affective processes. In addition, the insular cortex is often associated with detecting behaviorally relevant stimuli and salience processing (Uddin, 2015), however, as discussed later in section 3.2, some ICNs (69-73) overlapping with the insular cortex were grouped into the IT subdomain instead of the salience subdomain (SA) due to differences in their spatial distribution and functional modularity with other ICNs.

The STG (see ICNs 74-75) contains the auditory cortex involved in hearing (i.e., auditory perception). The medial surface (BA 41 and 42) makes up the primary auditory cortex (also known as A1, Heschl’s gyrus, or the transverse temporal gyrus) which receives auditory information from the inner ear via the thalamus and performs the most basic level of sound processing (Purves et al., 2001a). The posterior STG (BA 22; see ICN 74) has long been referred to as Wernicke’s area (typically in the left hemisphere) and has been heavily studied for its role in language comprehension (both written and spoken; Javed et al., 2024).

#### 2.6.2. Higher Cognition Domain - Temporoparietal Subdomain

The TP includes five ICNs (see ICNs 76-80) primarily centered in the posterior temporal (BA 22) and IPLs (particularly the angular gyrus, BA 39), converging at the temporoparietal junction (TPJ), although some of the ICNs reveal weaker but notable activation in the precuneus and inferior frontal gyrus as well. The TPJ (see ICNs 76-78) and posterior temporal lobe (see ICNs 79-80) have been implicated in various cognitive functions, particularly language and social cognition (Bahnemann et al., 2010; Hertrich et al., 2020; Igelström & Graziano, 2017). Specifically, these regions appear to have an important role related to Theory of Mind functions (i.e., conceptualizing and distinguishing between the self and others; Bahnemann et al., 2010; Hertrich et al., 2020) as well as discourse comprehension (i.e., representing and interpreting semantic meaning; Lin et al., 2018). Although Theory of Mind and language networks are functionally distinct, they appear to be synchronized during rest and language comprehension (Paunov et al., 2019).

#### 2.6.3. Higher Cognition Domain - Frontal Subdomain

The FR is spatially large, containing 10 ICNs (see ICNs 81-90) spanning across regions of the frontal lobe, including the inferior frontal gyrus (BAs 44, 45, & 47; including ventrolateral prefrontal cortex, VLPFC), middle frontal area and dorsolateral prefrontal cortex (DLPFC; BA 46), orbitofrontal (BA 11) and frontopolar (BA 10) cortex, as well as the frontal eye field area (FEF; BA 8) and anterior SMA (BA 6). The frontal cortex has been studied extensively for its involvement in the most complex forms of cognition (e.g., memory, language, intelligence, and executive functions requiring working memory and attention, such as problem solving), with higher levels of complexity associated with more anterior regions (Levy, 2024; Mesulam, 2000; Thiebaut de Schotten et al., 2017). The inferior frontal gyrus (IFG; see ICNs 81-86) lies anterior to the precentral sulcus and superior to the lateral fissure and is divided cytoarchitecturally into pars opercularis (BA 44), pars triangularis (BA 45), and pars orbitalis (BA 47). The lateral portion of the IFG is often referred to functionally as VLPFC which is typically associated with decision making and emotion regulation (Krawczyk, 2002; Mitchell, 2011; Sakagami & Pan, 2007). BA 44 and BA 45 are often referred to as Broca’s area which has traditionally been considered a key node in the language network and has been heavily studied for its role in speech production (e.g., motor movements which allow for speech, sentence structure, fluidity, etc.), although it also contributes to speech comprehension (e.g., semantics and phonology; Stinnett et al., 2024). The middle frontal gyrus is superior to the IFG and contains BA 46, which is functionally referred to as DLPFC (see ICN 88). This region is associated with various cognitive control functions, including decision making and emotion regulation (Mitchell, 2011), working memory and attention (Curtis & D’Esposito, 2003), and speech and language processing (Hertrich et al., 2020, 2021). The orbitofrontal cortex (BA 11; see ICN 87) is the most inferior portion of the frontal cortex, resting directly above the eyes. This region is also involved in decision making (Rogers et al., 1999), particularly aspects of reward processing (Kringelbach, 2005) and processing and encoding new information (Frey & Petrides, 2000). The frontopolar cortex (BA 10; also known as rostral or anterior prefrontal cortex; see ICN 88) is the most anterior region of the frontal cortex, and as such is also believed to contribute to the most complex cognitive functions (e.g., relational reasoning, combining and integrating information, decision making, etc.) and have the greatest capacity for the abstraction of information (Kroger & Kim, 2022; Levy, 2024; Thiebaut de Schotten et al., 2017). The FEF (BA 8; see ICN 89) is primarily associated with the control of visual attention and eye movement (Krauzlis, 2013) and is a key structure in the dorsal attention network, which is believed to be responsible for top-down controlled attentional selection (Vossel et al., 2014). In addition to the sensorimotor functions of the SMA described previously, the SMA (BA 6; see ICN 90) contributes to various cognitive control functions including complex response organizing and task-switching (Nachev et al., 2008), speech and language processing (Hertrich et al., 2016), and music performance (Tanaka & Kirino, 2017).

### 2.7. Triple Network Domain

There are 15 ICNs (see ICNs 91-105) within the triple network domain (TN). The ICNs in this domain display iconic patterns characteristic of triple network theory (Menon, 2011) commonly described in resting-state fMRI literature (Bremer et al., 2022; Lv et al., 2018; Menon, 2023). We have divided the ICNs within the TN domain into three subdomains, the central executive subdomain (CE), the default mode subdomain (DM), and the SA. With this domain label, we have made an exception to our previous nomenclature and chosen to uniquely include the word “network” due to its association with the triple network theory. Notably, the three large-scale brain networks defined in triple network theory are composed of many brain regions which naturally overlap with the domains we have previously described. Consistent with these spatial signatures, the ICNs in the TN domain also tend to be larger and overlap with multiple domains rather than fitting into a single domain. This overlap likely contributes to the strong modularity between the TN and other domains, particularly between the FR and CE (see Figure 3).

Although these networks are seemingly well-known, the structures included within them vary greatly across studies. In fact, Uddin and colleagues (2023) suggest that the brain networks encompassed in this domain have the least amount of consensus and consistency among all the large-scale brain networks discussed in the current paper. One reason for this inconsistency, as we will elaborate below, is that the spatial patterns within these networks may be better described as a continuum rather than a trichotomy (see Figure 4). Although resting-state fMRI literature traditionally perceives these three networks as functionally and spatially distinct (Menon, 2011, 2023), we observed that many of the ICNs in these networks have overlapping spatial distributions, and as shown in Figure 4, there appears to be a spatial gradient between the ICNs in the CE, DM, and SA (see Figure 4). That is, among the ICNs in the DM, some display greater spatial overlap with ICNs in the CE, while others display greater spatial overlap with ICNs in the SA, and some display minimal spatial overlap with either.

**Figure 4.**
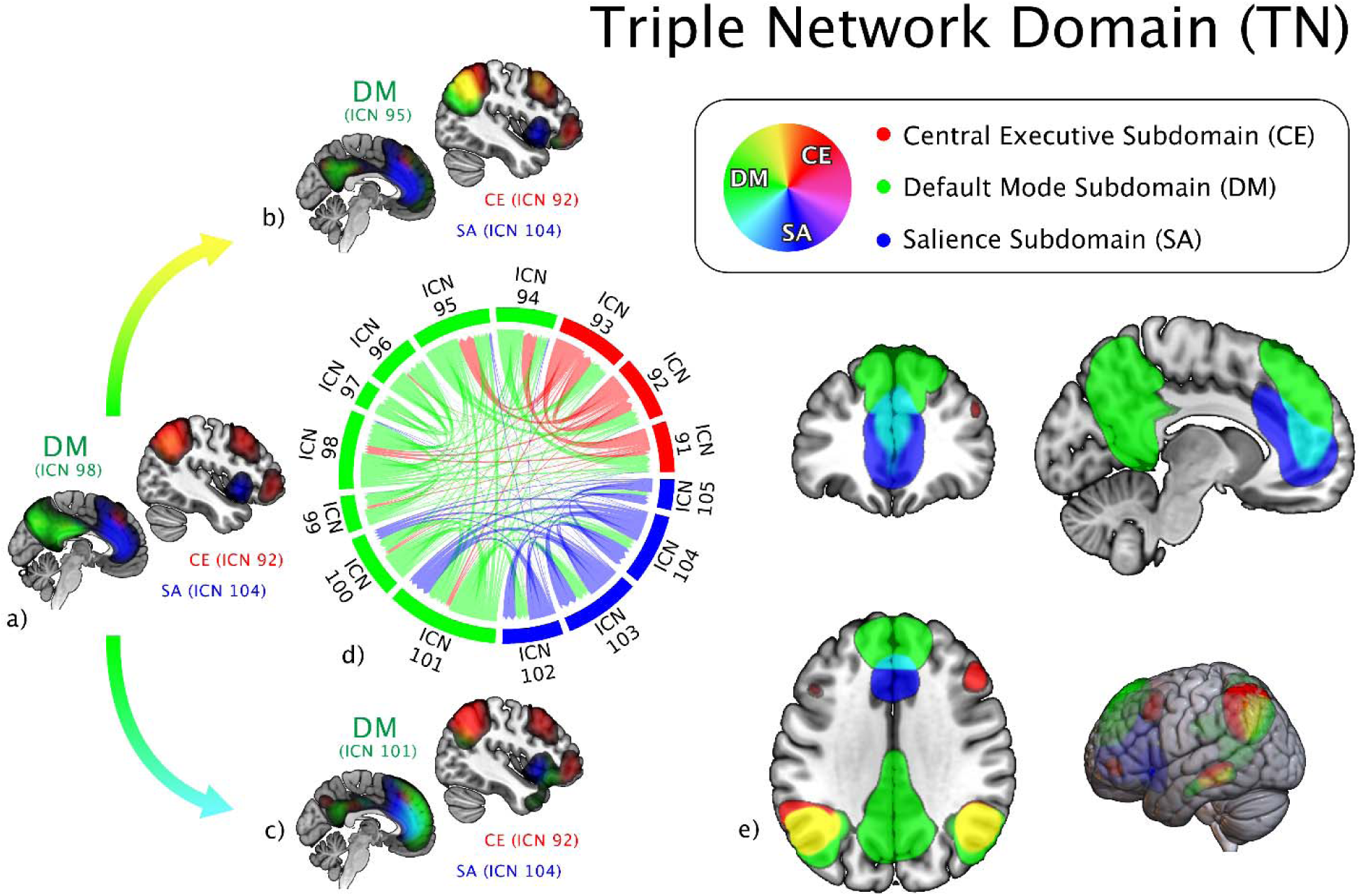
Spatial gradient of the triple network (TN) domain. Spatial maps for the TN intrinsic connectivity networks (ICNs) are overlayed on the MNI152 brain template above. The central executive (CE) subdomain ICNs are displayed in red, the default mode (DM) subdomain ICNs are displayed in green, and the salience (SA) subdomain ICNs are displayed in blue. Color is additive, with cyan representing the spatial overlap between the green DM and blue SA, and yellow representing the spatial overlap between the green DM and red CE. An apparent transition in the spatial maps of the DM ICNs are shown on the left, where a) DM ICN 98 displays relatively little overlap with CE and SA ICNs, b) DM ICN 95 displays greater overlap with CE ICN 92, and c) DM ICN 101 displays greater overlap with SA ICN 104. d) The proportion of significant voxels (z-score > 1.96) overlapping between all TN ICNs across subdomains is displayed. e) Spatial maps for all TN ICNs are overlayed on the MNI152 brain template.

#### 2.7.1. Triple Network Domain - Central Executive Subdomain

The CE includes three ICNs (see ICNs 91-93) which spatially correspond with the frontoparietal central executive network (CEN) described by Menon (2011). This subdomain consists primarily of DLPFC (BA 9 and 46) and parietal cortex (BAs 39, 40, and 7), but also overlaps with the FEF (BA 8) and middle temporal gyrus (MTG; BA 21). Our CE subdomain also appears to resemble networks referred to as the dorsal frontoparietal network (Corbetta & Shulman, 2002) and dorsal attention network (DAN; Vossel et al., 2014) which are active during goal-directed behavior. The CEN and DAN have both been characterized as a top-down attention control mechanism for maintaining and manipulating information in working memory during problem-solving and decision-making (Corbetta & Shulman, 2002; Menon, 2011; Vossel et al., 2014).

#### 2.7.2. Triple Network Domain - Default Mode Subdomain

The DM includes eight ICNs (see ICNs 94-101) spatially corresponding with the commonly known default mode network (DMN) frequently described in rsfMRI literature (Greicius et al., 2003; Menon, 2011; Raichle, 2015; Raichle et al., 2001). This subdomain consists primarily of the IPL (BA 39; see ICN 94-95, 98), posterior cingulate cortex (PCC; BA 23; see ICNs 94-98, 101), precuneus (BA 7; see ICNs 94- 99, 101), and medial prefrontal cortex (medial surface of BAs 8, 9, 10, & 32; see ICNs 94-95, 100-101), with some ICNs overlapping with the MTG (BA 21; see ICNs 94-95), superior/middle frontal gyri (lateral surface of BAs 8 and 9; see ICNs 94-95, 99-100), parahippocampal gyrus (BA 27; see ICNs 96-97), and retrosplenial cortex (BAs 26, 29, and 30; see ICNs 96-98). Although there are many brain regions associated with the DMN, previous investigations of dynamic connectivity have suggested that different sets of regions contribute to the DMN at different time points, which may result in different spatial patterns being identified for the DMN across different static analyses (Iraji, Fu, et al., 2019). This may account for some of the inconsistency in how the DMN is defined across studies. As indicated by its name, the DMN is typically observed to be less active during stimulus-driven cognitive tasks, with an increase in activity during rest (Menon, 2011; Raichle, 2015). However, it has been suggested that the DMN should not be viewed as a “task-negative” network because it can be active during goal-directed cognition depending on the task (Spreng, 2012). In other words, the DMN appears to be suppressed when attention is focused on external stimuli (i.e., while the SA or CE are active, although during some tasks these networks can be coupled with the DMN; Spreng, 2012) and becomes active when attending to internal thought processes such as self-referential judgments and self-other distinctions (Theory of Mind), episodic and autobiographical memory, language comprehension and semantic processing, and mind wandering (Menon, 2023).

#### 2.7.3. Triple Network Domain - Salience Subdomain

The SA includes four ICNs (see ICNs 102-105) spatially corresponding with the salience network (SN) described by Menon (2011) and Uddin (2015). This subdomain consists of the anterior cingulate cortex (ACC; BA 24 and 32; see ICNs 102-105) and anterior insular cortex (AIC; see ICNs 103-105). The SN has spatial overlap with the ventral frontoparietal network (Corbetta & Shulman, 2002) and the ventral attention network (Vossel et al., 2014), although the ICNs in our SA subdomain do not spatially overlap with parietal regions. The SN also overlaps with the cingulo-opercular network (Dosenbach et al., 2008), although the cingulo-opercular network also includes the anterior prefrontal cortex and thalamus. The SN has been characterized as a bottom-up attention processing mechanism for detecting unexpected but relevant stimuli and triggering a shift in attention (Corbetta & Shulman, 2002; Menon, 2011; Vossel et al., 2014). Vossel and colleagues (2014) described the relationship between dorsal and ventral attention networks (corresponding with CE and SA in our taxonomy) as dynamic interactions which enable flexible shifts in attention.

## 3. A Data-Driven Multi-Scale Functional Brain Atlas

### 3.1. Towards a Universal Taxonomy of Functional Brain Networks

Uddin and colleagues (2019) attempted to address the inconsistent nomenclature in network neuroscience by proposing a standardized taxonomy comprised of six large-scale brain networks named using anatomical terminology. We commend their efforts, although as the Workgroup for Harmonized Taxonomy of Networks (WHATNET) has acknowledged (Uddin et al., 2023), much confusion still exists. Presently, we have extended upon their work by employing a whole-brain multi-scale approach which is not limited to large-scale functional networks. Specifically, we have organized a data-driven functional brain template into multiple scales by grouping 105 individual ICNs into seven domains which can be further delineated into 14 subdomains. Many of the domains in our organization are comparable to those described by Uddin and colleagues (2019). Specifically, Uddin and colleagues’ (2019) occipital network (ON) corresponds with our visual domain, although notably, we include ICNs extending into the temporal lobe (e.g., the fusiform gyrus) which have high spatial overlap and functional modularity (see Figure 3) with the more traditionally “visual” occipital ICNs. Uddin and colleagues’ (2019) pericentral network (PN) corresponds with our sensorimotor domain although we have divided the ICNs corresponding with the STG and regions of the primary auditory cortex into our insular-temporal subdomain within the higher cognition domain. Uddin and colleagues’ (2019) dorsal frontoparietal network (D-FPN) and lateral frontoparietal network (L-FPN) both appear to correspond with our central executive subdomain, which might be described as a combined frontoparietal network, consisting of the parietal lobe (mostly inferior, but not exclusively), the FEF, and DLPFC. Together these two networks described by Uddin and colleagues (2019) would also draw from various ICNs within the frontal and temporoparietal subdomains within our higher cognition domain. Notably, our CE subdomain highlights potential overlap both spatially and functionally between the CEN and DAN described in rs-fMRI literature and suggests that these networks may not be as distinctly different as the terminology might imply. Alternatively, our subdomain may capture a dynamic relationship between multiple distinct networks. Uddin and colleagues’ (2019) midcingulo-insular network (M-CIN) corresponds with our salience subdomain, although ours only includes the core structures of the AI and ACC, while theirs suggests additional less well characterized areas such as the IPL, right TPJ, lateral prefrontal cortex, and various subcortical structures. Lastly, Uddin and colleagues’ (2019) medial frontoparietal network (M-FPN) corresponds with our default mode subdomain.

Importantly, our domains extend beyond the six large-scale networks defined by Uddin and colleagues (2019), which acknowledge but largely omit cerebellar and subcortical structures. Including and highlighting cerebellar and subcortical structures in our atlas is an important step towards overcoming cortico-centric bias (Parvizi, 2009; Saban & Gabay, 2023) which we hope will contribute to and expand our understanding of understudied areas of the brain as these regions are incorporated into network models and theories across the fields of neuroscience. Perhaps more importantly, our atlas consists of networks at different scales and addresses inconsistencies in methods, data, and subject variability both between and within studies by establishing a universal reference space. The labeling and organization we have applied to Iraji and colleagues’ (2023) multi-scale template has enabled it to evolve into a powerful functional brain atlas which can be used outside of the NeuroMark framework in other data-driven ICA approaches. For example, our atlas can be used with auto-labeler software (freely available in GIFT: http://trendscenter.org/software/gift; Salman et al., 2022) to label new networks in future studies using blind ICA. Alternatively, our atlas can also be used in combination with the recently released Network Correspondence Toolbox for comparison with widely used whole-brain parcellations that are not derived using ICA (Kong et al., 2024; available for download here: https://github.com/rubykong/cbig_network_correspondence). Furthermore, the utility of this atlas is not limited to fMRI, as it can be used to inform the labeling of sources obtained from other modalities such as positron emission tomography (PET).

### 3.2. Insights on the Integrative Nature of the Functional Brain

We have organized our functional brain atlas by grouping individual ICNs into domains and subdomains with the intention of providing a heuristic structure to aid in analysis by organizing results so that they can be more easily interpreted within existing frameworks in the field of neuroscience. However, the distinct boundaries of these functional domains are an illusion created by existing frameworks in the field of neuroscience; this is apparent in the strong patterns of modularity between domains and subdomains in the FNC matrix (see **Figure 3**). The grouping of ICNs into their assigned domains was based on how their associated brain regions are characterized in the literature as well as their functional and spatial modularity with other ICNs. The spatial similarity and FNC of ICNs typically display modular patterns across the defined domains, however, as demonstrated in **Figure 3**, there are some ICNs which display modular patterns across many domains (e.g., the insular ICNs 69-73), and others which appear to have relatively low modularity with nearly all domains (e.g., the frontopolar ICN 88).

This observation highlights an important distinction between dynamic functional sources and static anatomical brain regions. For this reason, there are some ICNs which have similar anatomical labels but have been grouped into different domains (see **Table 1**). For example, ICNs 69-73 which spatially overlap with the insula have all been grouped into the Insular-Temporal subdomain, while ICNs 104 and 105 which also spatially overlap with the insula have been grouped into the SA subdomain. Notably, ICNs 104 and 105 differ from 69-73 in that they also overlap with the ACC and demonstrate higher modularity with other ICNs in the SA subdomain and TN domain. Similarly, ICNs 88 and 89 which spatially overlap with the DLPFC and FEF have been grouped into the frontal subdomain, while others overlapping with these regions (ICNs 91-93) have been grouped into the CE subdomain. The divergent grouping of these ICNs into separate domains was again driven by spatial characteristics and unique patterns of modularity, where ICNs 91-93 also spatially overlapped with the IPL and displayed stronger functional modularity with other ICNs in the TN domain. As the field of neuroscience continues to develop, we continue to deepen our understanding of the highly integrative nature of the brain as well as the heterogeneity of function of individual brain regions (McCaffrey, 2015), and the boundaries we draw are likely to appear more and more arbitrary. Therefore, it is likely that our approach to organization will need to continuously update as frameworks within the field evolve.

Insights into the integrative nature of the brain are also evident in the unique spatial distributions of each individual ICN. As demonstrated in **Figure 2**, groups of ICNs which compose a functionally modular domain or subdomain may span across many different brain structures. However, it is also apparent that even across individual ICNs composing a given domain/subdomain, there are different proportions of spatial overlap with different anatomical brain regions. For example, from the spatial maps and chord chart for the IT in **Figure 2**, it can be observed that the ICNs span across the brain including regions in the temporal lobe (e.g., superior temporal lobe, middle temporal lobe, & temporal pole) and the insula, as well as subcortical regions such as the putamen. However, while overlap with these structures may be displayed across all ICNs in the IT, ICN 75 displays a high amount of overlap with the temporal lobe and little to no overlap with the insula. In contrast, ICN 71 displays the opposite pattern, with high spatial overlap with the insula and relatively little overlap with the temporal lobe. Despite the apparent differences in their spatial distributions, these two ICNs demonstrate a high level of functional modularity, suggesting that these ICNs are functionally similar (Bertolero et al., 2015; DeRamus et al., 2021; Ferrarini et al., 2009). This is consistent with the notion that specialized brain functions are accomplished through the recruitment of multiple brain structures throughout the brain, rather than being localized to a specific region (Just et al., 1999).

Perhaps the most effective example of blurred boundaries lies in the presence of a spatial gradient across the spatial maps of the ICNs within the TN, where the spatial maps of the DM ICNs appear to transition between the CE and SA subdomains (see **Figure 4**). This gradient is also reflected in the FNC matrix in **Figure 3a**, where unique patterns of modularity are displayed between subdomains within the TN. This spatial gradient is especially pronounced in ICNs 94 and 95 (see **Figure 4d**). We assigned these ICNs to the DM because of their spatial overlap with the precuneus, PCC, and medial prefrontal cortex (PFC), which are distinctive regions of the DMN (Buckner et al., 2008; Menon, 2011). However, these ICNs also have considerable overlap with posterior parietal cortex as well as DLPFC. Although the DMN is sometimes described as including posterior parietal cortex and the lateral surface of the superior/middle frontal gyri (Andrews-Hanna et al., 2014; Uddin et al., 2019), these regions serve as the primary structures of the CEN (Menon, 2011; Seeley et al., 2007). Consequently, the ICNs with the greatest overlap with ICNs 94 and 95 (i.e., ICNs 91-93) were assigned to the CE subdomain. From a quantitative perspective, ICNs 94 and 95 collectively have a great amount of spatial overlap (i.e., number of shared significant voxels) with ICNs categorized as DM, however, they have greater spatial overlap with the ICNs categorized as CE (see the proportions of green DM and red CE overlap in ICNs 94 and 95 in **Figure 4d**). However, in contrast to ICNs 94 and 95, the ICNs categorized as CE (ICNs 91-93) have little or no overlap with the precuneus, PCC, and medial PFC. Furthermore, these structures are not traditionally included in the CEN, therefore, we determined that assigning ICNs 94 and 95 to the DM subdomain would achieve greater consistency with existing literature. However, it is clear from their heterogenous spatial distributions, that these ICNs do not fit neatly into either category (i.e., CEN or DMN), but rather highlight the spatial overlap between both. The presence of this spatial gradient provides insight into the unique relationship between the three networks and may supply further evidence of the switching mechanism between them (see Menon, 2023). Future investigation into the nature of the enigmatic relationship between the functional and spatial signatures of these networks is warranted.

## 4. Conclusions

Selecting an atlas is an important methodological decision and we acknowledge that there are many atlases, and each has different advantages or disadvantages (Revell 2022). However, the atlas presented here provides several important benefits due to its whole-brain data-driven nature and the high replicability of its ICNs across individuals, datasets, and studies. We anticipate that the nomenclature and accompanying atlas presented here will provide a framework which will facilitate clear and insightful interpretations of results in many future functional brain imaging studies utilizing the NeuroMark templates. Furthermore, we anticipate that the terminology utilized within this template will influence the labels assigned to future templates and studies using data-driven approaches and may contribute more broadly to the field of neuroscience by informing the development of a universal taxonomy – or the “Brodmann Areas” of functional neuroimaging. The standardization of such an atlas would undoubtedly make functional neuroimaging techniques more accessible and enhance its utility and impact in cognitive and affective neuroscience and related fields.

## Acknowledgments

Funding Information: This work was supported by the National Institutes of Health grant number R01MH123610 (to VDC) and 5R01MH119251 (to AI) and National Science Foundation grant number 2112455 (to VDC). LQU is supported by R21HD111805 from NICHD and U01DA050987 from NIDA. In addition, KMJ received support from the Georgia State University Second Century Initiative (2CI) Doctoral Fellowship.

We would like to thank Emma H. Jensen for assistance with the creation of Figures 1, 3, and 4.

## Footnotes

1 Brodmann delineated the 52 brain areas based on “objective” cytoarchitectonic criteria, although his approach is not purely objective as it relies on the subjective judgments of the rater. Some have criticized the Brodmann Areas for lacking observer independency, reproducibility, and objectivity (Zilles & Amunts, 2010).

2 For each dataset the subject-specific FNC was calculated after standard preprocessing steps (see Du et al., 2020; Iraji et al., 2023) with a Pearson correlation between the time courses of the 105 subject-specific ICNs estimated for each subject followed by conversion to z-scores and averaging.

