## Supplementary Materials for "Addressing Inconsistency in Functional Neuroimaging: A Replicable Data-Driven Multi-Scale Functional Atlas for Canonical Brain Networks"

**Supplementary Materials S1**. **Individual Spatial Maps for 105 ICNs**

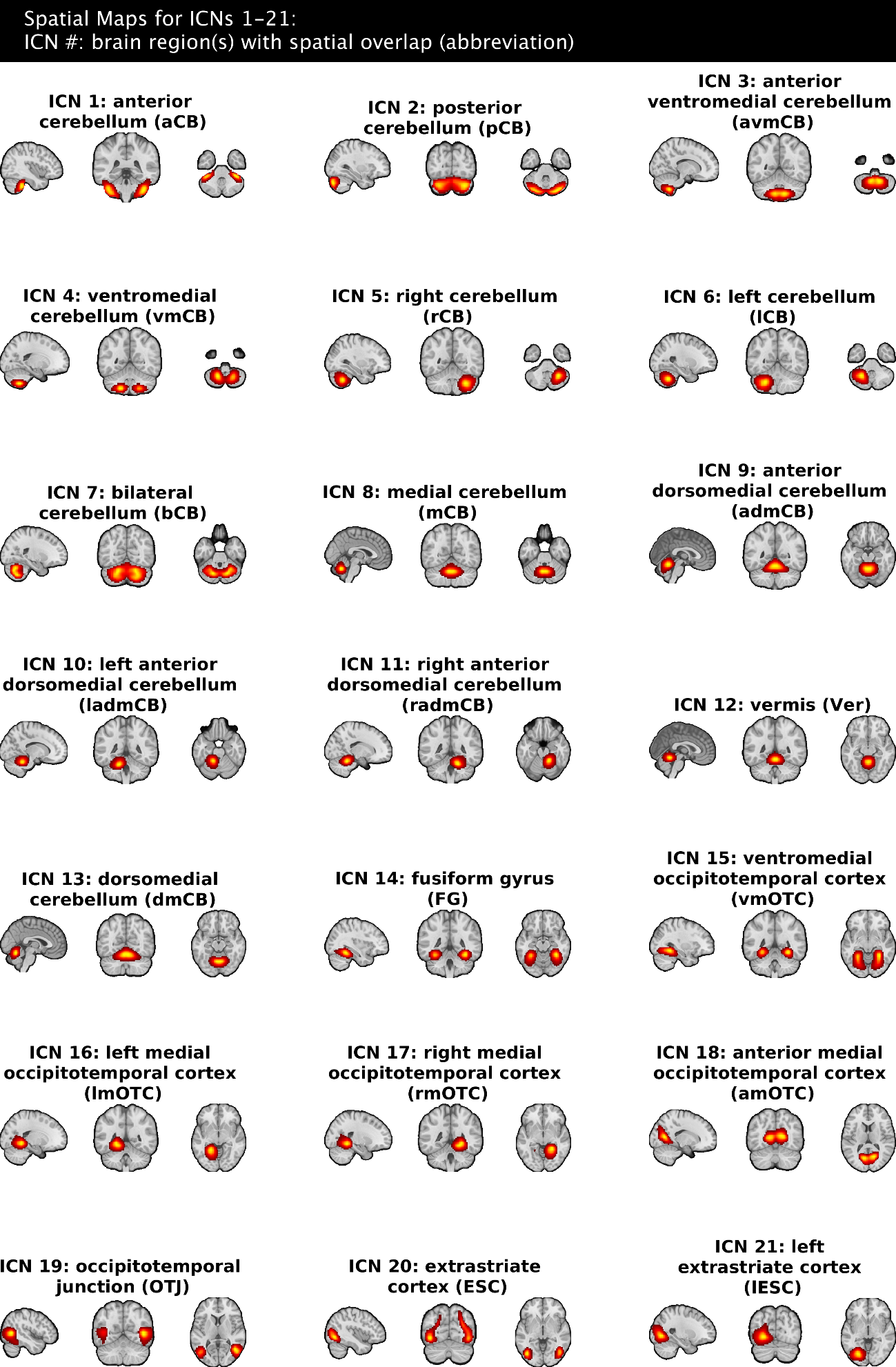

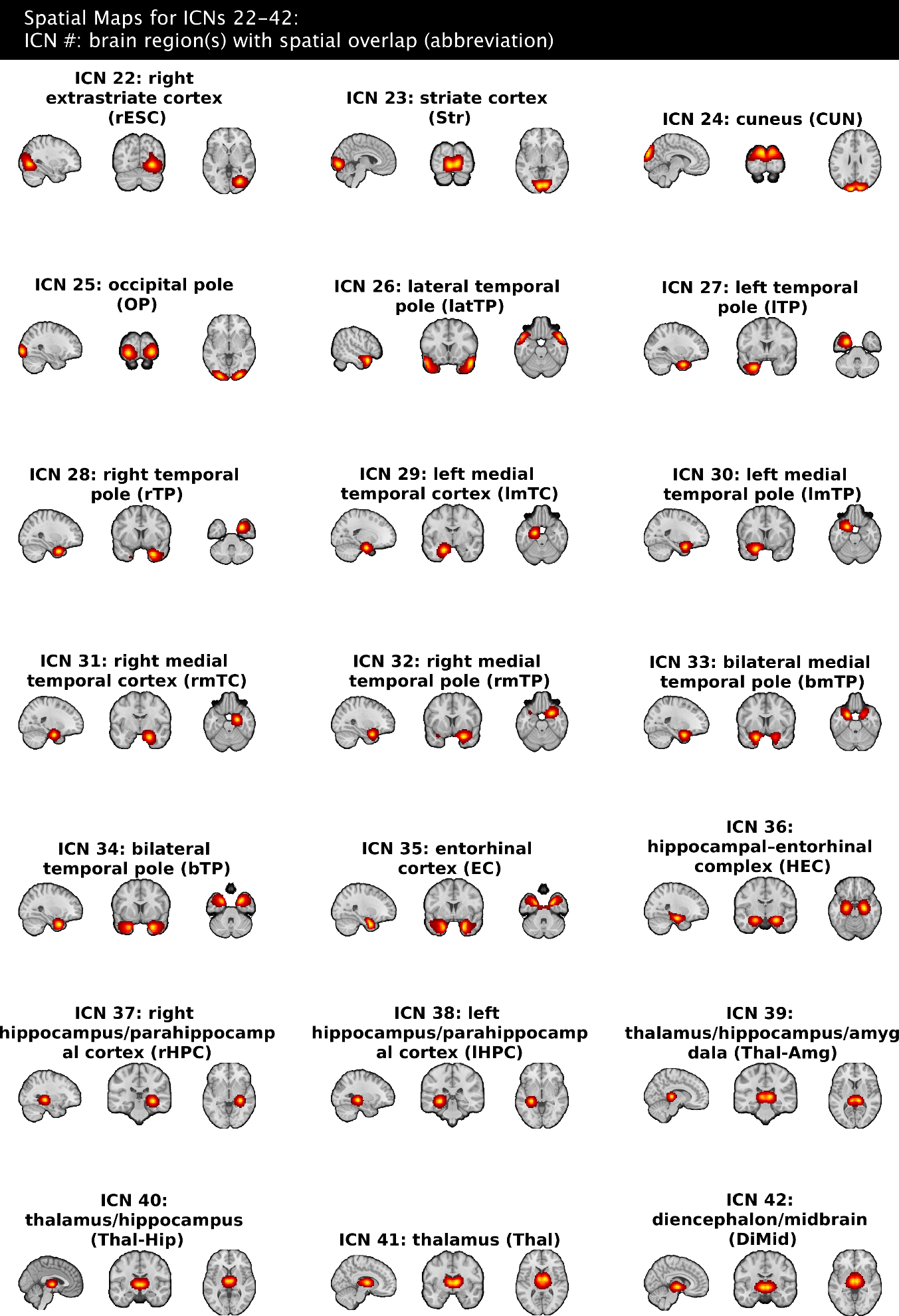

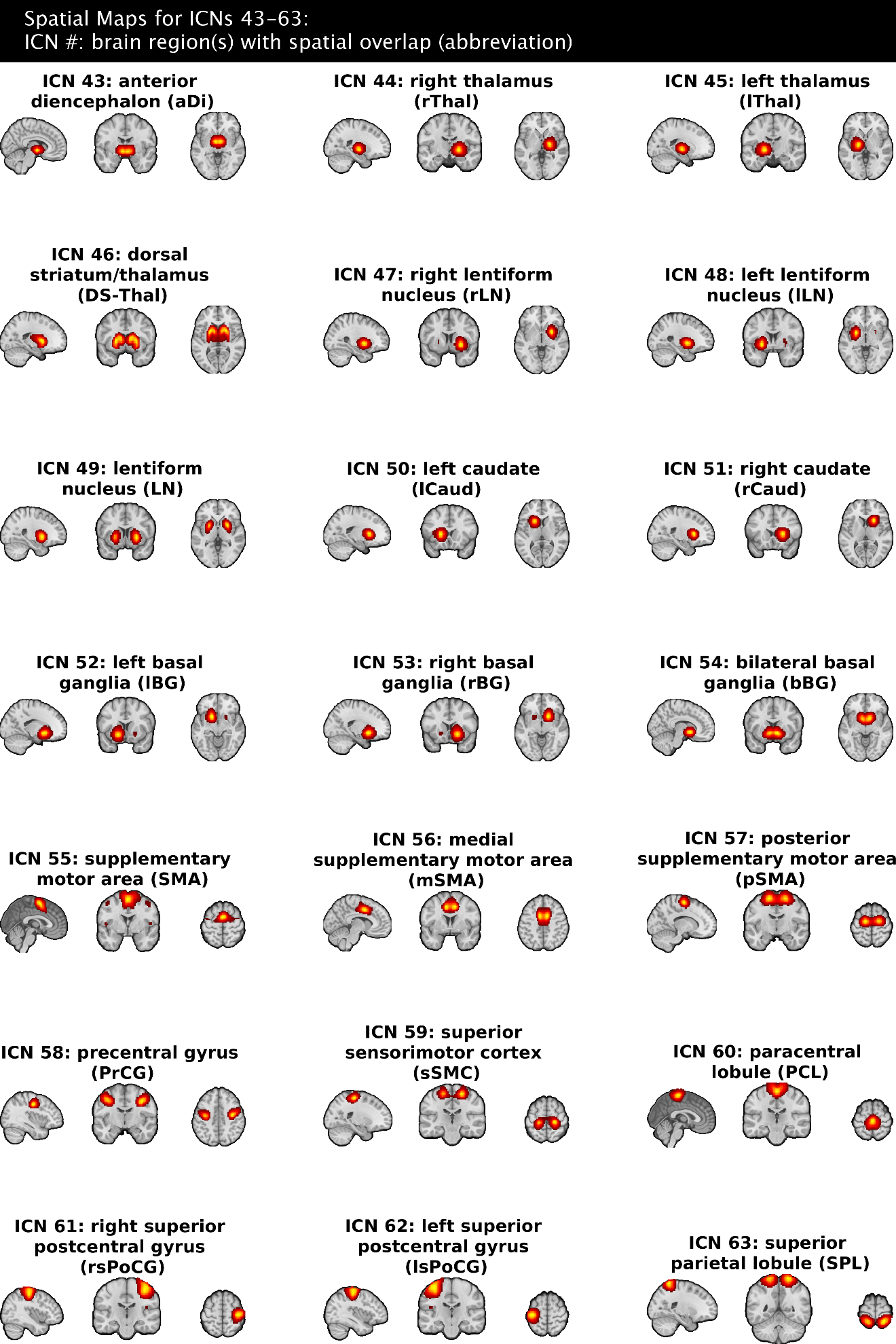

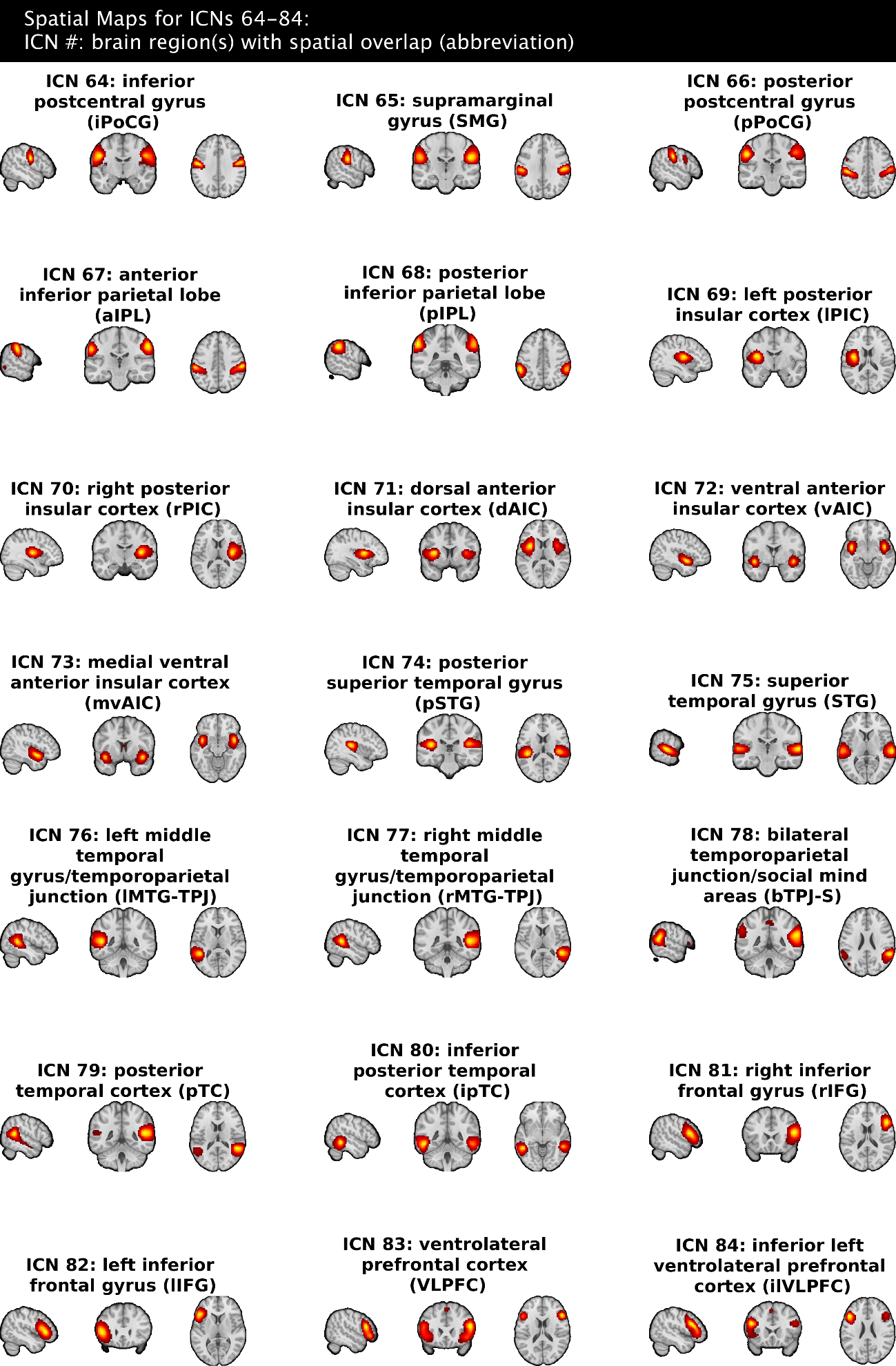

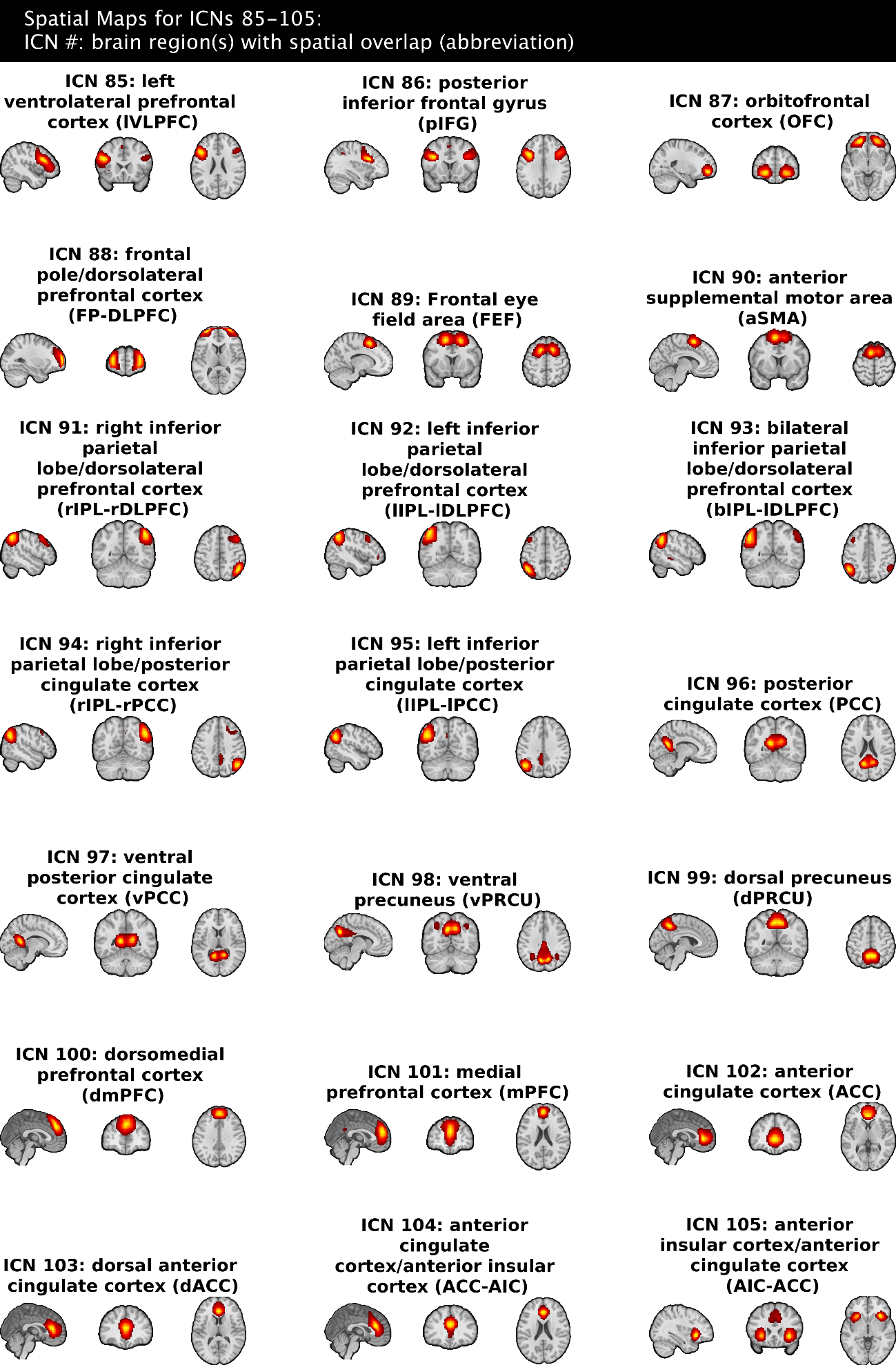

**Figure S1** | Spatial maps for each of the 105 intrinsic connectivity networks (ICNs) from the NeuroMark 2.2 multi-scale template are plotted above. Descriptions have also been provided for each ICN, characterizing them based on the anatomical and/or functional brain region(s) which they spatially overlap with. All spatial maps have been thresholded to display ICN voxels with a z-score > 3.

**Supplementary Materials S2**. **AAL Region Labels Table**

| Index | Structure |
| --- | --- |
| 1 | Precentral_L |
| 2 | Precentral_R |
| 3 | Frontal_Sup_L |
| 4 | Frontal_Sup_R |
| 5 | Frontal_Sup_Orb_L |
| 6 | Frontal_Sup_Orb_R |
| 7 | Frontal_Mid_L |
| 8 | Frontal_Mid_R |
| 9 | Frontal_Mid_Orb_L |
| 10 | Frontal_Mid_Orb_R |
| 11 | Frontal_Inf_Oper_L |
| 12 | Frontal_Inf_Oper_R |
| 13 | Frontal_Inf_Tri_L |
| 14 | Frontal_Inf_Tri_R |
| 15 | Frontal_Inf_Orb_L |
| 16 | Frontal_Inf_Orb_R |
| 17 | Rolandic_Oper_L |
| 18 | Rolandic_Oper_R |
| 19 | Supp_Motor_Area_L |
| 20 | Supp_Motor_Area_R |
| 21 | Olfactory_L |
| 22 | Olfactory_R |
| 23 | Frontal_Sup_Medial_L |
| 24 | Frontal_Sup_Medial_R |
| 25 | Frontal_Med_Orb_L |
| 26 | Frontal_Med_Orb_R |
| 27 | Rectus_L |
| 28 | Rectus_R |
| 29 | Insula_L |
| 30 | Insula_R |
| 31 | Cingulum_Ant_L |
| 32 | Cingulum_Ant_R |
| 33 | Cingulum_Mid_L |
| 34 | Cingulum_Mid_R |
| 35 | Cingulum_Post_L |
| 36 | Cingulum_Post_R |
| 37 | Hippocampus_L |
| 38 | Hippocampus_R |
| 39 | ParaHippocampal_L |
| 40 | ParaHippocampal_R |
| 41 | Amygdala_L |
| 42 | Amygdala_R |
| 43 | Calcarine_L |
| 44 | Calcarine_R |
| 45 | Cuneus_L |
| 46 | Cuneus_R |
| 47 | Lingual_L |
| 48 | Lingual_R |
| 49 | Occipital_Sup_L |
| 50 | Occipital_Sup_R |
| 51 | Occipital_Mid_L |
| 52 | Occipital_Mid_R |
| 53 | Occipital_Inf_L |
| 54 | Occipital_Inf_R |
| 55 | Fusiform_L |
| 56 | Fusiform_R |
| 57 | Postcentral_L |
| 58 | Postcentral_R |
| 59 | Parietal_Sup_L |
| 60 | Parietal_Sup_R |
| 61 | Parietal_Inf_L |
| 62 | Parietal_Inf_R |
| 63 | SupraMarginal_L |
| 64 | SupraMarginal_R |
| 65 | Angular_L |
| 66 | Angular_R |
| 67 | Precuneus_L |
| 68 | Precuneus_R |
| 69 | Paracentral_Lobule_L |
| 70 | Paracentral_Lobule_R |
| 71 | Caudate_L |
| 72 | Caudate_R |
| 73 | Putamen_L |
| 74 | Putamen_R |
| 75 | Pallidum_L |
| 76 | Pallidum_R |
| 77 | Thalamus_L |
| 78 | Thalamus_R |
| 79 | Heschl_L |
| 80 | Heschl_R |
| 81 | Temporal_Sup_L |
| 82 | Temporal_Sup_R |
| 83 | Temporal_Pole_Sup_L |
| 84 | Temporal_Pole_Sup_R |
| 85 | Temporal_Mid_L |
| 86 | Temporal_Mid_R |
| 87 | Temporal_Pole_Mid_L |
| 88 | Temporal_Pole_Mid_R |
| 89 | Temporal_Inf_L |
| 90 | Temporal_Inf_R |
| 91 | Cerebelum_Crus1_L |
| 92 | Cerebelum_Crus1_R |
| 93 | Cerebelum_Crus2_L |
| 94 | Cerebelum_Crus2_R |
| 95 | Cerebelum_3_L |
| 96 | Cerebelum_3_R |
| 97 | Cerebelum_4_5_L |
| 98 | Cerebelum_4_5_R |
| 99 | Cerebelum_6_L |
| 100 | Cerebelum_6_R |
| 101 | Cerebelum_7b_L |
| 102 | Cerebelum_7b_R |
| 103 | Cerebelum_8_L |
| 104 | Cerebelum_8_R |
| 105 | Cerebelum_9_L |
| 106 | Cerebelum_9_R |
| 107 | Cerebelum_10_L |
| 108 | Cerebelum_10_R |
| 109 | Vermis_1_2 |
| 110 | Vermis_3 |
| 111 | Vermis_4_5 |
| 112 | Vermis_6 |
| 113 | Vermis_7 |
| 114 | Vermis_8 |
| 115 | Vermis_9 |
| 116 | Vermis_10 |

**Table S2** | For the reader’s convenience, anatomical labels corresponding with the 116 brain regions in the Automated Anatomical Atlas (AAL; Tzourio-Mazoyer et al., 2002) are listed in the table above. These are intended to accompany the chord plots in Figure 2 and Figure S2.

**Supplementary Materials S3**. **Domain-Subdomain Overlap with Brodmann Areas**

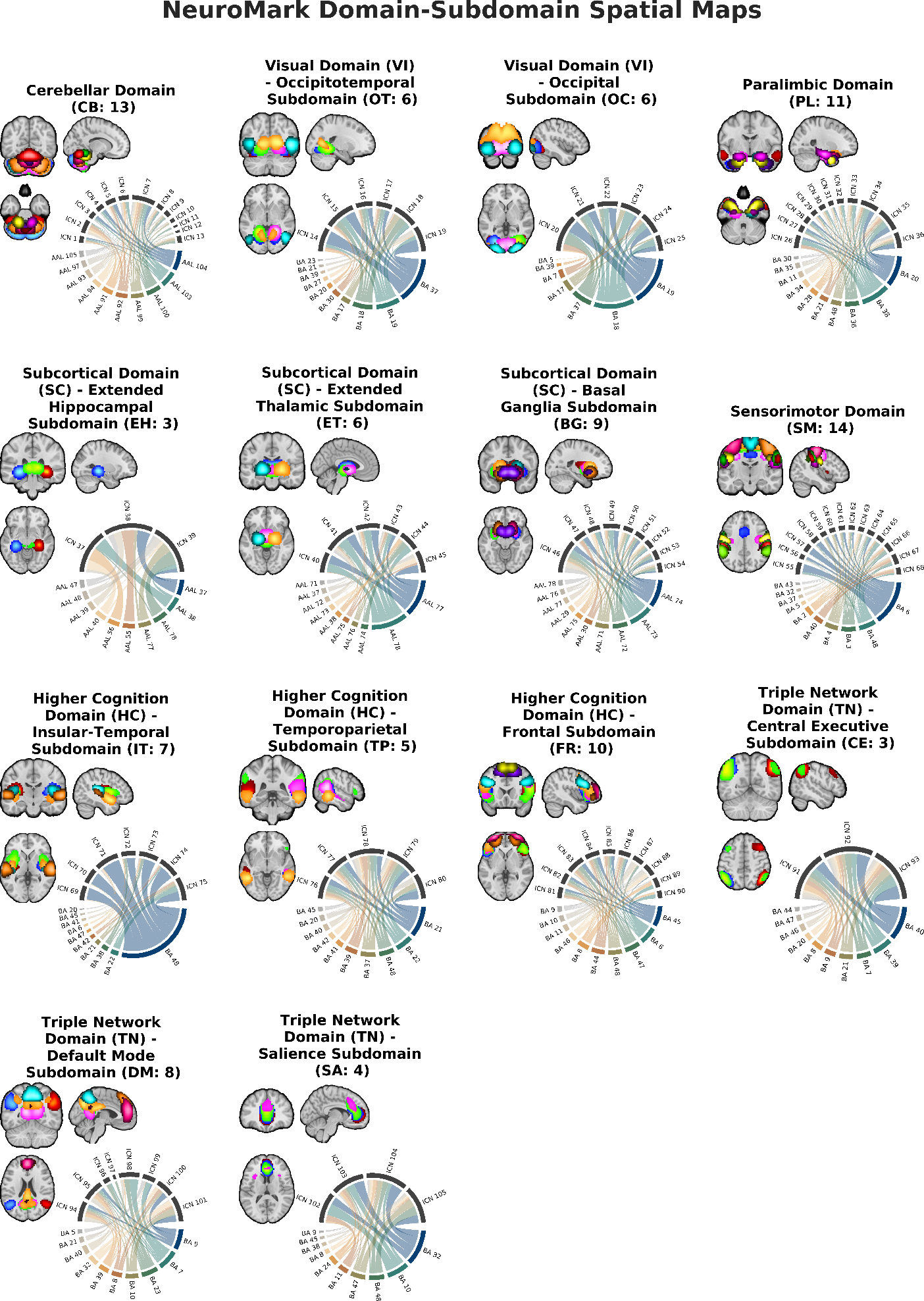

**Figure S3 |** Overlayed spatial maps of the 105 intrinsic connectivity networks (ICNs) from the NeuroMark 2.2 multi-scale template are plotted above for each of the seven domains and 14 subdomains: cerebellar (CB), visual-occipitotemporal (VI-OT), visual-occipital (VI-OC), paralimbic (PL), subcortical-extended hippocampal (SC-EH), subcortical-extended thalamic (SC-ET), subcortical-basal ganglia (SC-BG), sensorimotor (SM), higher cognition-insular temporal (HC-IT), higher cognition-temporoparietal (HC-TP), higher cognition-frontal (HC-FR), triple network-central executive (TN-CE), triple network-default mode (TN-DM), and triple network-salience (TN-SA). All spatial maps have been thresholded to display ICN voxels with a z-score > 3. In addition, chord charts are plotted for each subdomain, displaying the proportion of significant (z-score > 1.96) voxels overlapping with the top 10 most contributing Brodmann Areas (BA). Overlap with the top 10 most contributing regions from the automated anatomical atlas (AAL) are shown for Cerebellar and Subcortical domains, as these regions are excluded from Brodmann’s atlas.
